# An Atlas of Linkage Disequilibrium Across Species

**DOI:** 10.1101/2024.09.24.614726

**Authors:** Tian-Neng Zhu, Xing Huang, Meng-yuan Yang, Guo-An Qi, Qi-Xin Zhang, Feng Lin, Wenjing Zhang, Zhe Zhang, Xin Jin, Hou-Feng Zheng, Hai-Ming Xu, Shizhou Yu, Guo-Bo Chen

**Author notes:** Correspondence: Guo-Bo Chen, Shizhou Yu.

## Abstract

Linkage disequilibrium (LD) is a key metric that characterizes populations in flux. To reach a genomic scale LD illustration, which has a substantial computational cost of *𝒪*(*nm*^2^), we introduce a framework with two novel algorithms for LD estimation: X-LD, with a time complexity of *𝒪*(*n*^2^*m*) suitable for small sample sizes (*n <* 10^4^); X-LDR, a stochastic algorithm with a time complexity of *𝒪*(*nmB*) for biobank-scale data (*B* iterations); *n* the sample size, and *m* the number of SNPs. These methods can refine the entire genome into high-resolution LD grids, such as more than 9 million grids for UK Biobank samples (∼4.2 million SNPs). The efficient resolution for genome-wide LD leads to intriguing biological discoveries. **I)** High-resolution LD illustrations revealed how the pericentromeric regions and the HLA region lead to intense and extended LD patterns. **II)** Two universal LD patterns, identified as Norm I and Norm II patterns, provide insights on the evolutionary history of populations and can also highlight genomic regions of deviation, such as chromosomes 6 and 11 or ncRNA regions. **III)** The results of our innovative LD decay method aligned with the LD decay scores of 59.5 for Europeans, 60.2 for East Asians, and 33.2 for Africans; correspondingly, the length of the LD was approximately 2.85 Mb, 2.18 Mb, and 1.58 Mb for these three ethnicities. Rare or imputed variants universally increased LD. **IV)** An unprecedented LD atlas for 25 reference populations contoured interspecies diversity in terms of their Norm I and Norm II LD patterns, highlighting the impact of refined population structure, quality of reference genomes, and uncovered a profound *status quo* of these populations. The algorithms have been implemented in C++ and are freely available (https://github.com/gc5k/gear2).

## 1 Introduction

Linkage disequilibrium (LD) measures the non-random association between loci and encompasses a population that is in flux (1). LD decays by generation and is expected to be maintained in a short range, while extended LD can exist intrachromosomally and even interchromosomally (2; 3; 4). So far, LD patterns have often been investigated within a limited range (≤ 1,000kb), and global LD, which is the mean LD of *m* SNPs for *n* individuals, is rarely explored due to its computational time cost of *𝒪*(*nm*^2^) even with state-of-the-art methods (5; 6; 7; 8). In this study, we are able to investigate complex LD patterns in 29 publicly available datasets using our novel algorithms that reduce computational complexity from *𝒪*(*nm*^2^) **i)** to *𝒪*(*n*^2^*m*) for small to modest sample size (*n <* 10^4^), denoted X-LD (9); **ii)** to *𝒪*(*nmB*) for biobank-scale datasets such as UKB (*n <* 10^6^), denoted X-LDR.

Given our technical innovation, we carry out an unprecedented expedition, which has long been considered computationally prohibited, to draft an atlas of LD across species. We first explore how LD patterns are shaped throughout the human genome, such as by chromosomal length, allele frequency bins, imputation, and pericentromeric regions, which have recently unfolded after the release of the human reference genome T2T-CHM13 (10). We report interesting results as follows. **I)** For a random mating population, a longer chromosome leads to a much weaker average LD for markers, as often found in the human genome (11), and we call it Norm I pattern; in contrast, the Norm I pattern is distorted in many of the 25 species studied. **II)** Interchromosomal LD is ubiquitously observed and often proportional to the product of their chromosomal eigenvalues – an approximation of population strucutre; we call it Norm II pattern. **III)** Enriched rare or imputed variants lead to much extended LD. **IV)** Annotation-orientated LD patterns indicate that the LD strength of ncRNA is exceptionally higher than expected, as consistently found in Chinese populations and UKB populations. **V)** Through adjustment for population structure, the atlas can provide information on how LD has been shaped on each chromosome throughout their respective history in various species.

## 2 Overview of the study

### 2.1 Datasets

The study surveyed human and nonhuman samples (**Table 1**). Human samples included 1KG samples, unrelated White British of UKB (*n* = 278, 781; denoted as UKBW hereafter), unrelated Black British of UKB (*n* = 5, 057; UKBB) and multiethnicity samples of UKB (*n* = 487, 365; UKBA). For the UKB datasets, backbone SNPs were drawn from the UKB chipped SNPs, while rare SNPs were drawn mainly from the imputed SNPs. In addition, two Han Chinese cohorts were included: CONVERGE (*n* = 10, 640 Chinese women) (12; 13), and Westlake Biobank cohort (*n* = 4, 480), where the SNP call was performed using two different reference genomes: GRCh37 (denoted as WBBC hereafter) and T2T-CHM13 (WBBC-T2T hereafter). For human samples, excluding 1KG, the QC metrics for the inclusion of SNPs were: missing rate *<* 0.01 and minor allele frequencies (MAF) *>* 0.01 (no QC based on MAF was performed during the investigation of rare variants) for human samples. The other 25 datasets, denoted ***RefPop***, covered a depth of species, and their generic QC metrics for inclusion were autosomal SNPs only with missing call rates *<* 0.2, and MAF *>* 0.05. We extracted 4,257,224 common SNPs from CONVERGE and WBBC, and then the same set of SNPs was extracted from UKBW and UKBB; these four refined datasets were called CONVERGE-c, WBBC-c, UKBW-c, and UKBB-c, respectively.

**Table 1:**
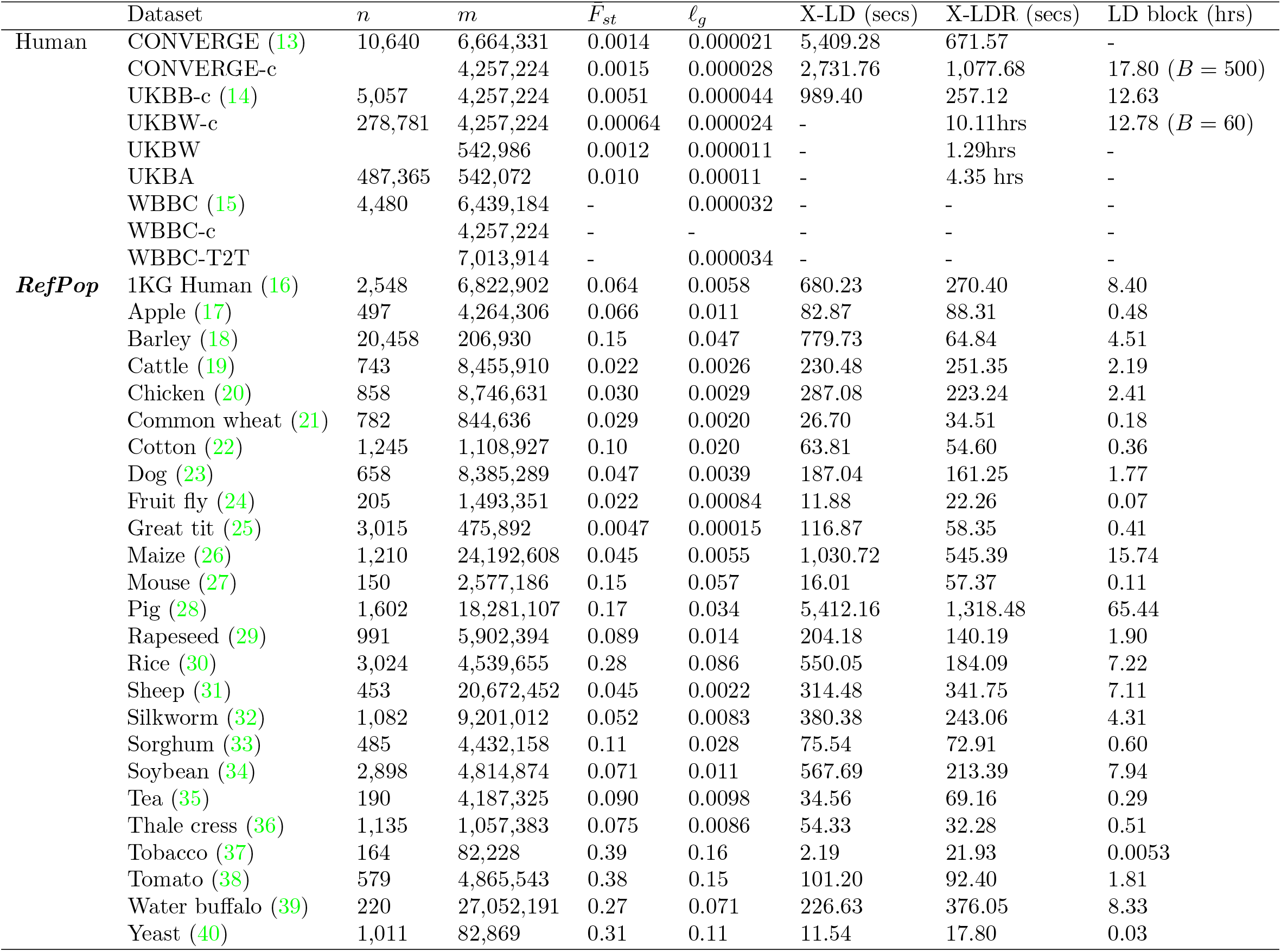
Performance of X-LD and X-LDR on various datasets. The genome-wide *F*_*st*_ is realized as 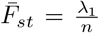, in which *λ*_1_ is the top eigenvalue of the dataset. The last three columns record the time for estimating 𝓁_*g*_ with XLD, XLDR, and for partitioning the whole genome into 1, 000 × 1, 000 SNP pairs for each genome (X-LDR for UKBW-c and X-LD for the other datasets). The computational time was measured on 80 threads.

### 2.2 Sketch of the analysis framework

#### 2.2.1 Efficient estimation for LD

For a standardized genotypic matrix **X** of *n* unrelated individuals and *m* SNPs (*n* ≪ *m*), we coin a transitive triplet

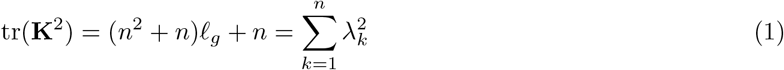

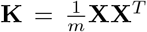, the genetic relationship matrix (GRM), *λ*_*k*_ the *k*^th^ eigenvalue of **K**, and 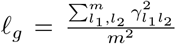 which 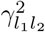 the LD metric (squared Pearson correlation) of the two subscripted SNPs. The relationship of 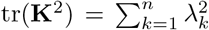 is well known in matrix algebra, and using Isserlis’s theorem, we can create the triplet (41; 42). As eigenvalues capture population structure (43), so after some rearrangement we have

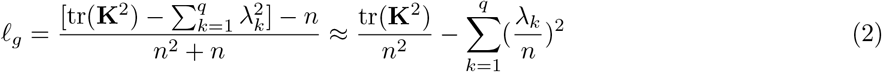

in which 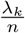 reflects population structure that is absorbed in each eigenvalue. When *q*≠ 0, 𝓁_*g*_ is consequently adjusted by population structure, and we call this adjustment technique *peeling* hereafter.

Conventionally, 𝓁_*g*_ has a computational time cost *𝒪*(*nm*^2^), which drains computational resources given substantially large *n* and *m* (*m* ≫ *n*) (7; 8). In this study, tr(**K**^2^) is estimated by our proposed methods X-LD or X-LDR, the computational cost for 𝓁_*g*_ is reduced from *𝒪*(*nm*^2^) to either *𝒪*(*n*^2^*m*) or *𝒪*(*nmB*), in which *B* is the iteration round for X-LDR (see **Methods**).

#### 2.2.2 Two norms of LD patterns

As the analysis for 𝓁_*g*_ can be for chromosome *i* (𝓁_*i*_) and ever for chromosomes *i* and *j* (𝓁_*ij*_), we propose a pair of regression models, LD-dReg and LD-eReg, to illucidate two global patterns of LD.

##### LD-dReg analysis for Norm I pattern

In a random mating population, 𝓁_*i*_ is reciprocal to its chromosomal length – the number of SNPs (*x*_*i*_) on chromosome *i*, and we have a linear model

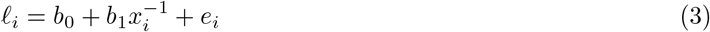

which is called LD decay regression (LD-dReg) (9). For a random-mating population, *b*_0_ approximates the population structure and *b*_1_ the mean LD score for an averaged LD block. If LD can be precisely estimated after the *peeling* technique, the 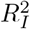 (determination coefficient) of LD-dReg will increase and justify the optimal eigenvalues for **Equation 2**. Pearson correlation *ρ*_1_ is used to reflect a possible negative correlation between 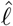 and 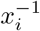, albeit pathological, as will be seen in ***RefPop***. Notably, the Norm I pattern stands on two factors: i) the LD decay equation *D*_*t*_ = *D*_0_(1 − *c*)^*t*^, in which *D*_*t*_ (*D*_0_) is LD at the *t*^*th*^ (0) generation and *c* the recombination for the pair of loci in question; ii) the relative uniform distribution of the recombination spots along a genome.

##### LD-eReg analysis for Norm II pattern

In terms of interchromosomal LD 𝓁_*ij*_, **Equation 1** can be modified analogously to be 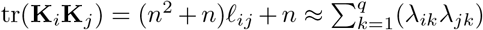 the *k*^th^ eigenvalue of **K**_*i*_. We can efficiently estimate 𝓁_*ij*_ using the following equation.

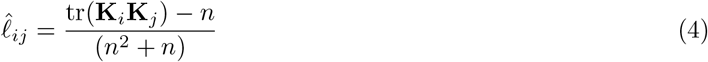

Little is known about interchromosomal LD, but is often assumed to be close to zero. We aim to regress 𝓁_*ij*_ with 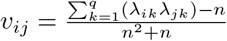, which we refer to as LD eigenvalue regression (LD-eReg).

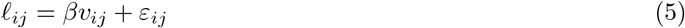

in which 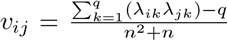. If there is population structure, the Norm II pattern will be observed inter-chromosomally, and 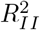 and *ρ*_2_ can be defined accordingly for LD-eReg (see **Methods**).

As the Norm I and Norm II patterns are reciprocal in elucidating LD, we will use them as guidelines to contour LD for various species. Detailed characteristics of LD will be explored depending on the source of the data.

## 3 Proof-of-principle validation

### 3.1 Reconciliation for LD estimators

Between LD estimation, such as the one implemented in PLINK (“–r2”), and that of our methods, there was a predictable quantity of 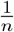. As X-LD has previously been validated against PLINK in the 1KG Human cohorts (9), we only compared the consistently estimated 𝓁_*i*_ and 𝓁_*ij*_ in X-LD with those of PLINK for yeast (*n*=1,011 and *m*=82,869), which was computationally feasible for PLINK (**Figure S1**). In ***RefPop***, X-LD and X-LDR showed high consistency for the estimations of 𝓁_*i*_ and 𝓁_*ij*_ in the 25 datasets (**Figure S2**).

### 3.2 Computational efficiency test

For the estimation of 𝓁_*i*_ (or 𝓁_*g*_) or 𝓁_*ij*_, we used X-LD when *n <* 10^4^ and X-LDR when *n >* 10^4^. Benchmark comparisons for X-LD and X-LDR were made in the 29 aforementioned public datasets evaluated on 80 computational threads (**Table 1**). In terms of time cost, to complete CONVERGE it took 21.53 hours by X-LD but 1.27 hours with the more efficient X-LDR only. In the text below, we chose X-LD for ***RefPop***, WBBC, UKBB, and CONVERGE, and X-LDR for UKBW and UKBA because X-LD became computationally infeasible. It took 0.57 hours to estimate genomic LD (𝓁_*g*_) of UKBA under 80 threads, while theoretically it would have taken 3,110.5 CPU hours for conventional methods (7; 8).

Of UKBW-c, we refined its genomic LD 𝓁_*g*_ into 9,114,315 LD blocks, each of which had 1, 000 × 1, 000 SNP pairs or a block size of 0.7 × 0.7Mb^2^ approximately (**Figure 1B**). As tested, X-LDR took 12.7 hours (*B* = 60 and 80 threads) to estimate the LD for these blocks. The high LD blocks were within the genomic neighborhood, along the chromosome as shown, but extremely high 𝓁_*i*_ blocks were found around the human leukocyte antigen (HLA) region on chromosome 6 and the pericentromeric regions of chromosomes 6 and 11. Theoretically, it should be possible to produce such an LD illustration using conventional methods with 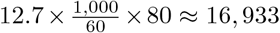 CPU hours, but its logistic and storage costs would have been far more complicated. We also refined genomic LD for CONVERGE-c and UKBB **Figure S3**, and for other populations, their computational time can be found in the last column of **Table 1**.

**Figure 1:**
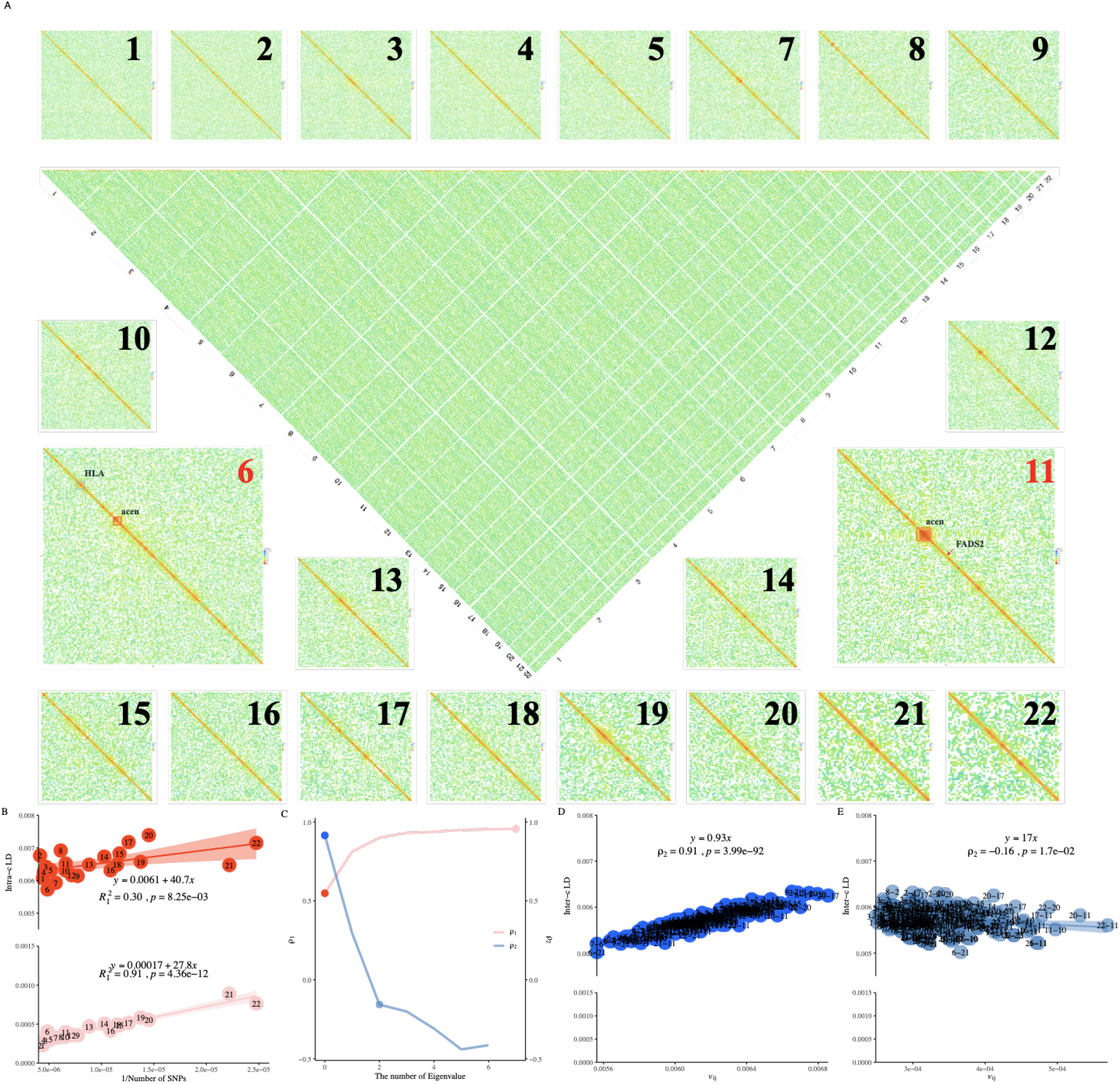
Illustration for the proposed methods in UKBiobank and 1KG human. **A**, Illustration of 9,114,315 LD blocks for UKBW-c (*n* = 278, 781, and *m* = 4, 257, 224). X-LDR took about 12.78 hrs (80 threads) to complete the computing. **B**, 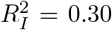 (without adjustment) and 0.91 after adjustment for the top 8 eigenvalues of 1KG human samples. The fitted LD-dReg models are printed. **C**, Dynamics for the Norm I and the Norm II (**D, E**) patterns after *peeling* for 1KG human samples. The detailed Norm I pattern with or without *peeling*, red and light red points, can be found in (**B**), and the Norm II pattern, blue and light blue points, in **D** and **E. D**, Norm II analysis for 1KG population using LD-eReg. **E**, Norm II analysis for 1KG population using LD-eReg after *peeling* for the top 2 eigenvalues of 1KG human samples. The fitted LD-eReg models are printed in **D** and **E**.

### 3.3 Norms I and II patterns in human 1KG population

The Norm I pattern was demonstrated in the 1KG population, which consisted of 26 cohorts of various ethnicities. After *peeling*, LD-dReg had its 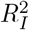 progressively increased from 0.30 to 0.91 (red and pink point in **Figure 1B, C**), and *ρ*_*I*_ between 𝓁_*i*_ and 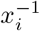 closely approached the ideal value of 1 (*ρ*_1_ = 0.91). Furthermore, as indicated by LD-dReg that 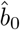 decreased from =0.0065 to 0.00057 and 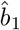 from 40.7 ± 13.9 to 27.8 ± 1.9, also indicated that *peeling* successfully reduced the inflation of LD caused by the population structure. As a negative control, we also estimated the LD-dReg model for each of the 26 cohorts without *peeling*, and, as none exhibited drastic inflation like the entire 1KG population, each had 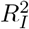 greater than 0.6 (**Figure S4**).

The Norm II pattern was also demonstrated in the 1KG population. For consistency, we used the top 8 eigenvalues of each chromosome in the LD-eReg. Due to the drastic admixture in the 1KG population, resulting in a strong population structure, 𝓁_*ij*_ fit very well with 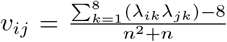 in LD-eReg, achieving 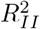 of almost 1.00 (**Figure 1C, D**), indicating the existence of interchromosomal LD but due to the population structure. Similarly, we calculated *ρ*_2_ in each of the 26 cohorts, none of which exceeded 0.4 (**Figure S5**).

As shown in **Figure 1C**, it was a reciprocal relationship between the Norm I and Norm II patterns in the 1KG population. After surrogating the population structure with eigenvalues, the Norm I pattern (*ρ*_1_) increased after gradually removing the population structure, while the Norm II pattern (*ρ*_2_) decreased.

## 4 LD atlas of the human genome

Our efficient LD estimation methods offer an opportunity to explore large-scale LD patterns for populations. As the human genome has much better annotated genomic information and other resources such as GWAS hits, we analyze the complex LD patterns for human cohorts. So, given the refined LD illustration (**Figure 1B**), we will explore more details of the characteristics of LD in humans, which will be a good benchmark to reflect the complex LD patterns in ***RefPop***.

### 4.1 Norm I pattern consistency across ethnicity

We estimated their respective 22 autosomal LD (𝓁_*i*_) for CONVERGE-c, WBBC-c, UKBW-c, and UKBB-c in their 4,257,224 shared SNPs. As expected, the Norm I pattern that 𝓁_*i*_ was reciprocally related to its chromosomal length was consistently observed in the four cohorts tested, and the order of 𝓁_*i*_ in different ethnic groups was ranked as follows: East Asians *>* Europeans *>* Africans (**Figure 2A**). East Asians and Europeans had their LD patterns resemble each other, and the African cohort exhibited a much smaller 𝓁_*i*_ because historical recombination events decay LD (44). Chromosomes 6 and 11 stood out for their much stronger 𝓁_*i*_. The upward deviation of chromosome 11 was driven by its more deeply sequenced pericentromeric region (46.06-59.41Mb) than other chromosomes (also see **Figure 1B**). In contrast, although there is known selection for *FADS2* on chromosome 11, a gene that is selected for a high animal fat content in Europeans and Chinese (45; 46), but no high LD found near *FADS2* (61.58-61.63Mb, **Figure 1B**) in neither UKBW nor CONVERGE-c **Figure S6**.

**Figure 2:**
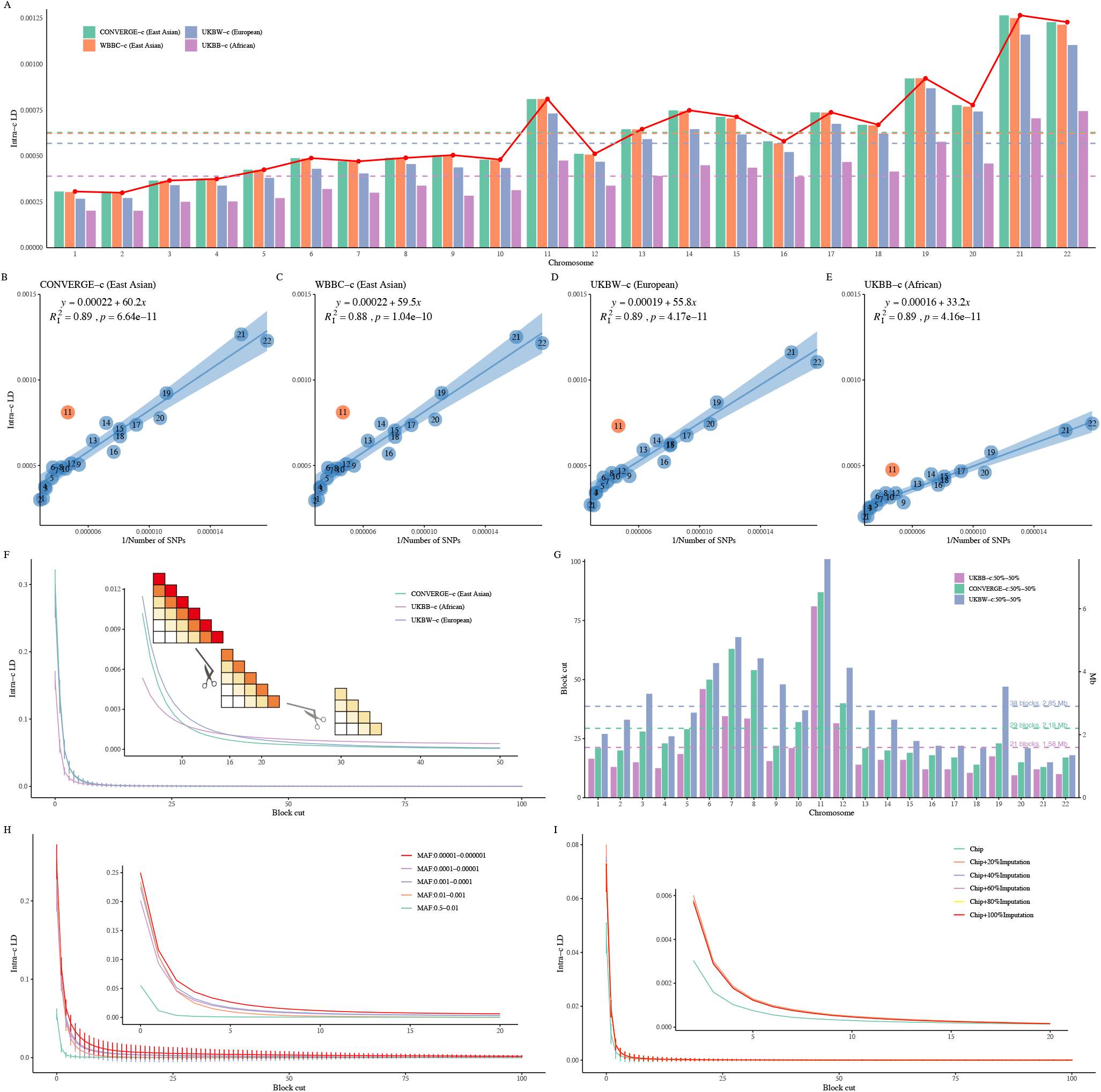
The consistent LD patterns in single-ethnicity human population. **A**, Average LD for each chromosome across four single-ethnicity populations; the red points represent the mean 𝓁_*i*_ across the four cohorts for each chromosome. The horizontal dashed line indicates the mean 𝓁_*i*_ for 22 autosomes for each color-matched cohort. **B-E**, The Norm I pattern analysis with LD-dReg. Chromosome 11 is found to be the most deviated chromosome in CONVERGE-c, WBBC-c, UKBW-c, and UKBB-c, respectively. The LD-dReg is displayed in each plot, and the quantity of the regression coefficient reflects mean LD score in each tested cohort. **F**, The LD decay plot is based on adjacent 100 LD blocks in CONVERGE-c, UKBB-c, and UKBW-c, respectively. A vertical line on each curve represents the standard deviation for 22 autosomes at each block cut. A cartoon is shown as subplot to illustrate how LD decay is performed. **G**, The run-off of LD blocks of the three cohorts on 22 autosomes. Each horizontal dashed line represents the mean LD decay blocks (axis at left) or distance (axis at right) for each cohort. **H**, The LD decay plot is based on adjacent 100 LD blocks across five MAF bins in UKBW. **I**, The LD decay plot is based on adjacent 100 LD blocks with increasing imputed SNPs in UKBW.

### 4.2 LD score and range estimation

We estimated the LD score using LD-dReg, the slope (*b*_1_) of which reflected the mean LD score of the high-LD blocks along the diagonal (9). First, these four cohorts produced a nearly identical Pearson correlation *ρ*_1_ of 0.94 and 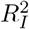 of 0.9 (**Figure 2B-E**); as observed, both CONVERGE-c and WBBC-c had nearly identical LD decay scores of 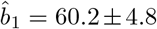 and 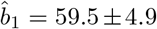, respectively, and UKBW-c had 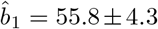. In contrast, UKBB-c had much smaller 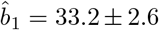 due to historical recombination events that decayed LD (**Table 2**). So, the magnitude of the LD score was also ordered as Asians ≥ Europeans *>* Africans.

Traditional LD decay methods typically analyze a limited range (generally up to 1,000kb), while we proposed an innovative LD decay analysis for extended LD based on high-resolution LD blocks. We partitioned the 4,257,224 shared SNPs into 906,230,164 blocks, each of which had a size of 75 × 75kb^2^ or 100 × 100 SNP pairs. Here we aggregated the results for all 22 autosomes by calculating the mean LD for each autosome at each block distance (**Figure 2F**). The results for each of the 22 autosomes individually are presented in **Figure S7**. When adjacent blocks were segmented at the top several blocks, the relative strength of LD between ethnicities remained consistent. However, when the distance increased to 16 blocks or more UKBB-c appeared to gradually exceed that of CONVERGE-c and UKBW-c, and UKBB-c had LD extended to 100 blocks (approximately 7.50Mb or 10,000 SNPs) or even further.

As the true LD decreased with physical distance, the remaining LD of UKBB-c was more plausibly due to population structure. We further elucidated the extension of LD by splitting each cohort into two even divisions and regressing their matched blocks against each other on each chromosome (**Figure S8**). For example, on chromosome 1 (**Figure S8A**), within CONVERGE-c, *R*^2^ between the two divisions diminished when the distance reached approximately 20 blocks (≈ 1.50Mb, approximately 2,000 SNPs); To determine a more precise block number at which true LD could be, we plotted a 95% confidence interval area of 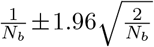, where *N*_*b*_ represents the remaining number of blocks after cutting (gray shadow in the subplot of **Figure S8A**). When the *R*^2^ bar was immersed in the shadow, it indicated that LD was running off and that LD in CONVERGE-c disappeared near the 21st block (≈ 1.58Mb, approximately 2100 SNPs) and near the 27th block for UKBW-c (≈ 2.03Mb, approximately 2,700 SNPs). In UKBB-c, due to certain population structure, *R*^2^ persisted even for up to 30 blocks (**Figure S8A**). To simply rule out the effect of population structure, we used the mean of the two crossover points where the LD of UKBB-c exceeds that of UKBW-c and CONVERGE-c as the number of LD blocks for the complete decay of LD in UKBB-c. We then derived the number of LD blocks for each chromosome across four single-ethnictiy populations (**Figure 2G**). By calculating the mean across the 22 chromosomes, we found that the average lengths of the LD block were approximately 1.58Mb for Africans, 2.18Mb for East Asians, and 2.85Mb for Europeans. A similar LD decay (≈ 2.78Mb) was observed in UKBW-c when it was divided into men and women (UKB field ID: 22001).

In general, the area under the fitted curves (**Figure S8G**) for each cohort was proportional to their estimated LD score of 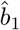 (**Figure 2B-E**). However, it seemed paradoxical that UKBW showed more extended LD (2.85Mb) but smaller 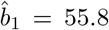, while CONVERGE-c showed less extended LD (2.18Mb) but greater 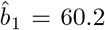. To further rule out the possible influence of MAF, we only used 1,783,915 SNPs that had MAF ≥ 0.05 and allelic difference ≤ 0.1 between UKBW and CONVERGE, but the results were almost identical (**Figure S9**).

#### 4.2.1 Influence of MAF in shaping LD pattern

Using the large sample size of UKBW, we investigated how MAF categories would lead to differential patterns of LD (47). We observed that 𝓁_*ij*_ increased for more rare variants in UKBW (**Figure S10**). To exclude any influence from imputation, we retained rare SNPs within the pre-imputation UKB dataset as a benchmark. Despite containing fewer variants, the pre-imputation UKB dataset exhibited similar LD patterns across MAF bins compared to the post-imputation dataset. We also applied LD block decay to each MAF bin (each block covering approximately 0.25Mb) and observed that rare SNPs showed stronger LD with a longer range (**Figure 2H, Figure S11**).

To investigate how imputation impacts LD, we extracted 733,367 chip SNPs from filtered UKB imputation SNP sets (10,203,936 SNPs, QC score ≥ 0.95). The remaining 9,470,569 SNPs were sampled from imputated SNPs. We gradually added 20% more imputed SNPs into the chip SNPs, resulting in six datasets with increasing numbers of SNPs. For analysis, we focused on 𝓁_*i*_ and divided each chromosome into equal size blocks (0.25 × 0.25Mb^2^) in each dataset. We applied LD-block decay analysis to these six datasets and observed that LD increased with the inclusion of more imputed SNPs (**Figure 2I, Figure S12**).

### 4.3 LD patterns shaped by structural and functional features

#### 4.3.1 High LD shaped by ncRNA

According to the functional annotation, 4,257,224 SNPs were classified into 9 functional categories (48). The Norm I pattern was consistently observed across different ethnicities when classifying the entire genome into functional categories as well. However, the categories of intronic ncRNA and exonic ncRNA stood out due to their significantly stronger LD (**Figure 3A**). We further validated this LD pattern using LD-dReg, where the LD scores 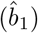 were 11.3 ± 10.65, 11.1 ± 10.48, 10.1 ± 9.13, and 7.51 ± 5.19 for CONVERGE-c, WBBC-c, UKBW-c, and UKBB-c, respectively, consistent with the trend of Asian *>* European *>* African (**Figure 3B-E**). It was also echoed by a previous study, but surveyed at a much smaller scale, that the regions around ncRNA had much higher LD (49).

**Figure 3:**
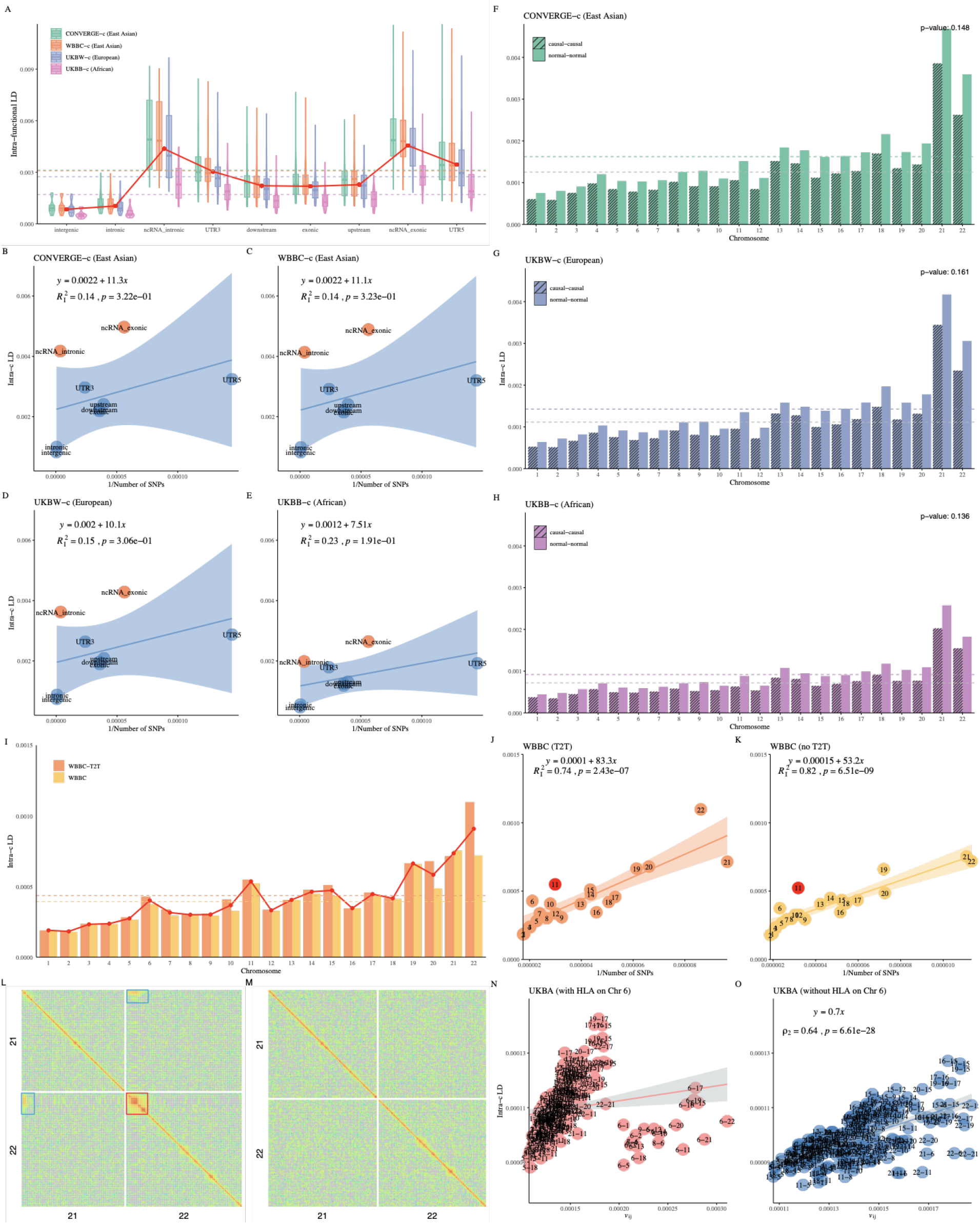
Influence of MAF (rare SNPs), chromosomal structure, and population structure on LD patterns in a single-ethnicity human population. **A**, Average LD for each functional region in CONVERGE-c, WBBC-c, UKBW-c, and UKBB-c, respectively. **B-E**, LD-dReg based on functional region in these four cohorts. **F-H**, Average LD for causal and non-causal variants (SNPs) of each chromosome in CONVERGE-c, UKBW-c, and UKBB-c, respectively. **I**, Average LD for each chromosome for WBBC and WBBC-T2T. **J, K**, LD-dReg analysis for WBBC and WBBC-T2T, respectively. **L**, High-resolution LD illustration of chromsome 21 and 22 for WBBC-T2T. The red box highlights the centromere, while the blue boxes indicate the regions of extremely high 𝓁_*ij*_. **M**, High-resolution LD illustration of chromosome 21 and 22 for CONVERGE. **N, O**, The Norm II pattern analysis with LD-eReg for UKBA (with and without the HLA region on chromosome 6).

#### 4.3.2 LD patterns of GWAS hits

One of the yet enigmatic parameters is LD between causal variants and SNPs. Because the exact number of causal variants segregating throughout the genome remains unknown, we chose to surrogate the distribution of causal variants from a recent fine-mapping result, which reported 7,209 causal segments for height in a GWAS meta-analysis of more than 5 million individuals (50). We extracted 745,748 “causal variants” from these causal segments, and then we shifted each segment by a distance equal to its length so that to create matched “not causal” segments and to extract 745,748 normal SNPs as controls. We then performed X-LD analysis in the CONVERGE-c, UKBW-c, and UKBB-c, respectively. Although causal and normal SNPs were found to have no significant differences in their estimated LDs that *p*=0.148, 0.161, and 0.136 in CONVERGE-c, UKBW-c, and UKBB-c, respectively, the causal SNPs actually had slightly lower LD than the control SNPs (**Figure 3F-H**).

#### 4.3.3 LD patterns shaped by centromeres

The human genome has recently been completely sequenced, and an obvious characteristic is dense satellite DNA sequences around the pericentromeric regions (51). We performed X-LD and LD-dReg on both WBBC and WBBC-T2T, the latter of which was assembled on T2T-CHM13, we found that almost each 𝓁_*i*_ of WBBC-T2T was higher than that of WBBC. In particular, chromosome 22 had the most significant increase, with 52.67% increase using T2T-CHM13 as a reference (WBBC-T2T) (**Figure 3I**). WBBC-T2T obviously had a higher LD score of 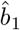 = 83.3 ± 10.9, while WBBC had 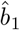 = 53.1 ± 5.6 (**Figure 3J, K**). The pericentromeric regions in WBBC-T2T had extremely high 𝓁_*i*_, especially on chromosome 22. Surprisingly, a distinct pattern of high 𝓁_*ij*_ was specifically identified around the pericentromeric regions between the acrocentric chromosomes, such as chromosome 21 and chromosome 22 (**Figure 3L**). According to chromosome band information on GRCh37, we have identified that 𝓁_*ij*_ originated from the heterochromatic regions and the pericentromeric regions located on the short arms of chromosomes 21 and 22. This observation implied the potential existence of conserved sequences, predominantly satellite sequences, in the vicinity of the centromere regions of various acrocentric chromosomes. As a negative control, we also estimated the LD blocks for CONVERGE (X-LDR, 22,294,504 blocks, 1, 000 × 1, 000 SNP pairs), but no such high LD pattern was found in CONVERGE (**Figure 3M**).

Furthermore, we extracted the pericentromeric regions of the 22 autosomes in WBBC-T2T and CONVERGE, respectively. To explore a clearer LD pattern around the centromere, we plotted the adjacent 20 LD blocks for chromosome 22 in WBBC-T2T (**Figure S13A**) and CONVERGE (**Figure S13C**). As the centrometric region was sequenced more completely in WBBC-T2T (**Figure S13B**, using T2T-CHM13 as the reference genome), WBBC-T2T exhibited a large LD block around the pericentromeric region, while CONVERGE did not due to the low density of SNPs around the pericentromeric regions (**Figure S13D**). Moreover, we also found that LD was higher in its flank regions than in its centrometric center, so if recombination events were relatively even around the pericentromeric region, it was consistent with the discovery that the centromeric centers were younger and the two flanks older (51).

#### 4.3.4 Distorted Norm II pattern by the HLA region

We found that the points between chromosome 6 and other chromosomes significantly deviated from its LD-eReg analysis in UKBA (**Figure 3N**). This deviation was driven by the HLA region on chromosome 6. When we removed the HLA region in UKBA and performed LD-eReg again (**Figure 3O**), the deviation of chromosome 6 disappeared as expected. We interpret this phenomenon as follows: the LD-eReg essentially inferred the correlation between 𝓁_*ij*_ and the product of *F*_*st*_ of a pair of chromosomes. If each chromosome has undergone genetic drift (even gentle polygenic selection) or is in a strong mixture (such as the 1KG population), it resulted in similar LD patterns on each chromosome, allowing 𝓁_*ij*_ to fit well with the production of *F*_*st*_ of chromosomes *i* and *j*. Factors such as pericentromeric regions and the HLA region can disturb chromosomal homogeneity, thus distorting LD-eReg. Of UKBA, chromosome 6 exhibited a significantly larger eigenvalue than other chromosomes, due to the influence of the HLA region, which accounted for the observed deviation of the results of LD-eReg. However, no deviation related to chromosome 6 was observed in the 1KG population (**Figure 1E**). UKB samples and CONVERGE were also included as references in the analysis below when necessary.

## 5 LD atlas across species: *RefPop*

Metaphorically, if a human population, such as UKBW, was a flat plane in which detailed elements distorted the LD pattern were likely to be elucidated, the 25 species in ***RefPop*** not only different from human populations but dramatically from each other (**Figure 4**). ***RefPop*** showed a substantial population structure (**Table 1**), the general correlation between 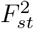 and 𝓁_*g*_ was 0.98 for ***RefPop***, indicating an influence due to their respective population structure. We created a novel Kilt plot that illustrates the high-resolution genomic LD for each species, and its computational time could be found in **Table 1**. Some species had strong diagonals, such as apple and cattle (**Figure 4B, D**), but some had unexpected elements along the diagonal, such as sorghum, and soybean (**Figure 4R, S**); there were sometimes extremely large LD block so visible that even ran along a chromosome, such as barley and cotton (**Figure 4C, G**), or even interchromosomally, such as tobacco and tomato (**Figure 4V, W**). Consequently, we aimed to sketch up an overview of ***RefPop*** in terms of their Norm I and Norm II patterns, and then to explore the details for certain species.

**Figure 4:**
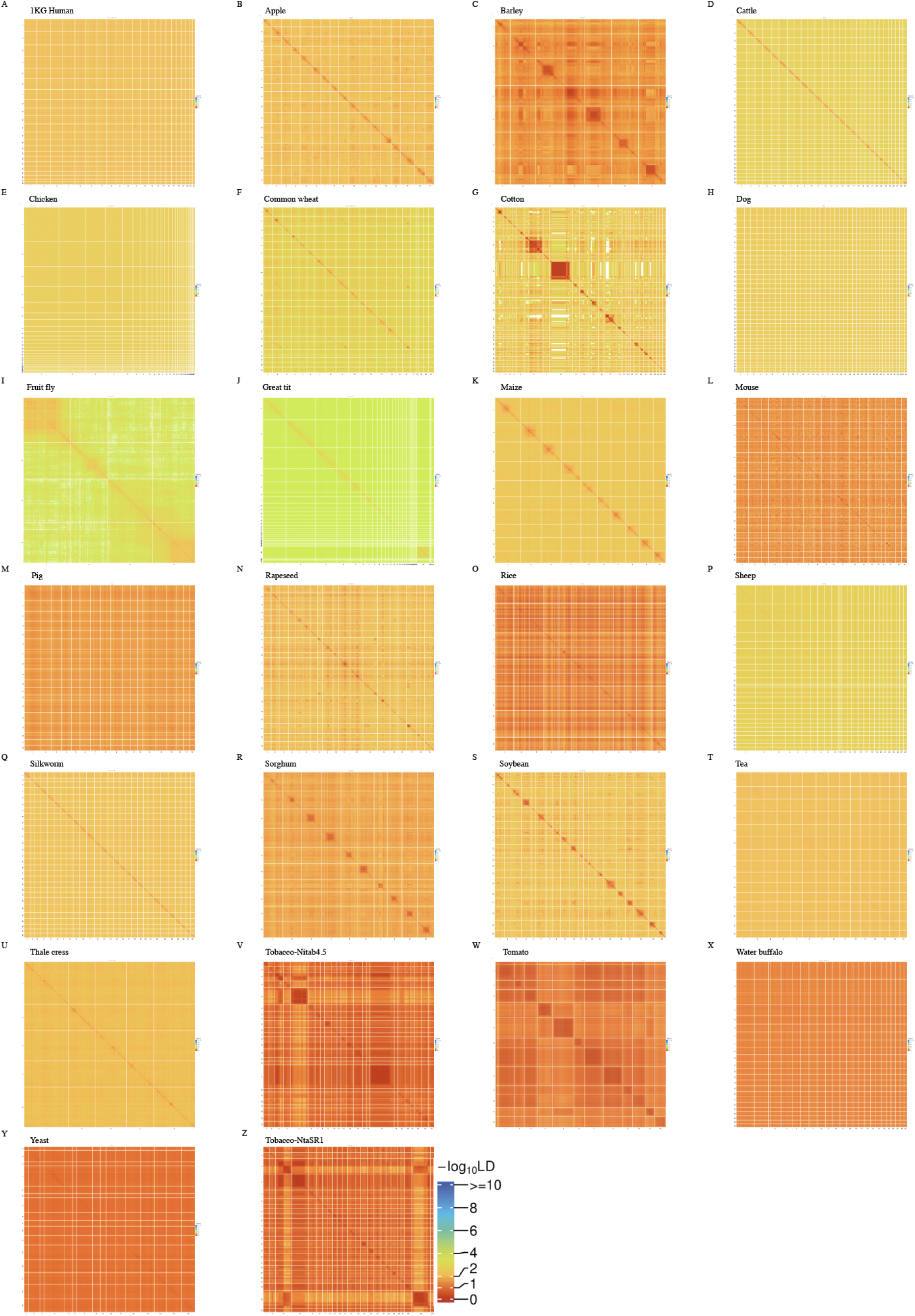
Kilt plot for 25 species in *RefPop*. **A-Y**, LD illustration for each species in ***RefPop***. The resolution of LD is at a unit of 1,000 × 1,000 SNP pairs. The color scheme of LD is on the same scale and comparable across species. **V** and **Z** are from the same set of SNPs but on the tobacco reference genomes Nitab4.5 (52) and NtaSR1 (37), respectively; the chromosome 17 of tobacco is unproportionally large in

### 5.1 Variation of the Norm I and Norm II patterns in *RefPop*

In LD-dReg analysis for ***RefPop***, the variation of 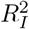 and *ρ*_1_ before and after *peeling* was displayed (**Figure S14**). Before *peeling* LD-dReg revealed a continuous distribution of *ρ*_1_, ranging from −0.77 (tomato) to 0.94 (UKBB), but increased significantly after *peeling*, indicating that the *peeling* method could extract the population structure and revealed a more natural pattern of LD (**Figure S15I**, *t* -test, *p* = 1.02 × 10^−6^). As indicated by colored lines, the human cohorts had a much higher *ρ*_1_ (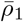=0.73 and 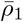=0.99 after *peeling* for the latter) than those nonhuman cohorts (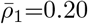 and 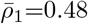, **Figure 5I**). Notably, the diverse 1KG human population could have its *ρ*_1_ increased from 0.36 to 0.97 after *peeling* of its top 16 eigenvalues. Some ***RefPop*** species, however, showed yet negative *ρ*_1_ after *peeling*, specifically, cattle (*ρ*_1_ = −0.64 and *ρ*_1_ = −0.86 after *peeling* for the latter), common wheat (*ρ*_1_ = −0.48 and *ρ*_1_ = −0.68), fruit fly (*ρ*_1_ = −0.58 and *ρ*_1_ = −0.53), mouse (*ρ*_1_ = 0.22 and *ρ*_1_ = −0.61), tobacco (*ρ*_1_ = −0.091 and *ρ*_1_ = −0.25), and tomato (*ρ*_1_ = −0.77 and *ρ*_1_ = −0.44), apart from the Norm I pattern. Therefore, the distortion of *ρ*_1_ may not only due to population structure. More detailed results of the Norm I results can be found in **Extended Data 1**.

**Figure 5:**
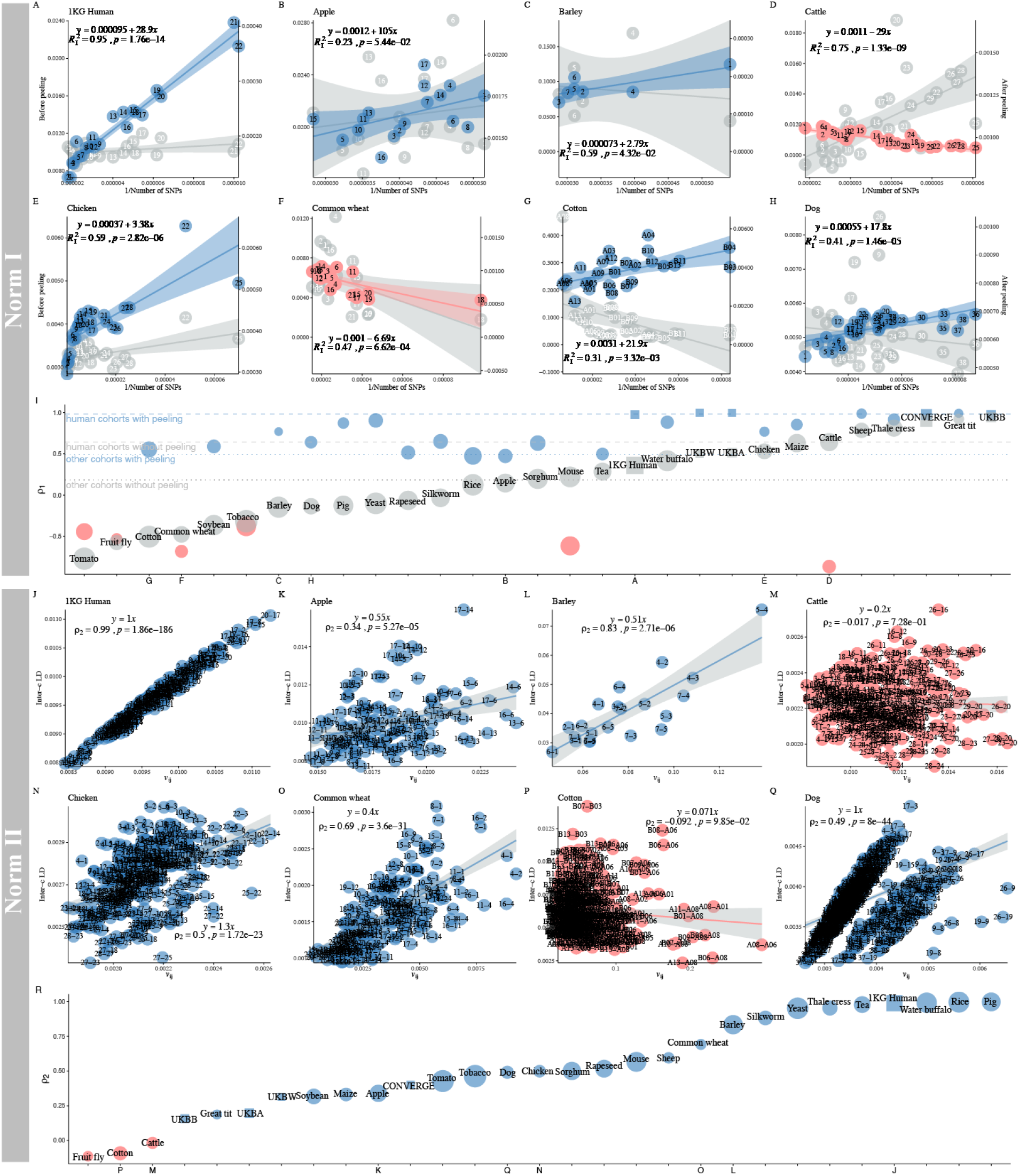
LD atlas on the Norm I and Norm II patterns in *RefPop*. **A-H**, The LD-dReg before and after *peeling* of eight representative species in ***RefPop*. I**, The *ρ*_1_ of the Norm I pattern in ***RefPop***, ordered by *ρ*_1_ before *peeling* (human populations are denoted by squares, while other populations are represented by circles). The size of each square or circle is proportional to the 𝓁_*i*_ of each population, with larger shapes indicating higher 𝓁_*i*_. The blue dot and dashed lines represent 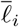 of the human cohorts (1KG human, UKBW, UKBB, and UKBA) before 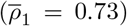 and after *peeling* 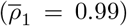, and the gray dot and dashed lines represent 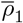 of the nonhuman species in ***RefPop*** (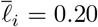 and 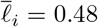). **J-Q**, The LD-eReg for the same eight species in ***RefPop*. R**, The *ρ*_2_ between 𝓁_*ij*_ and 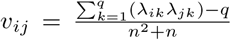 in ***RefPop***, ordered by *ρ*_2_ of LD-eReg. Human populations are represented by squares, while other populations as circles. The size of each square or circle is proportional to the 𝓁_*ij*_ of each dataset, with larger shapes indicating higher 𝓁_*ij*_.

Differing from the Norm I pattern, which was enhanced after peeling off the population structure, in their Norm II patterns (**Figure S16A-Y**), the populations with strong population structures corresponded to a good fit in their LD-eReg models (*ρ*_2_ ≈ 1), such as pig 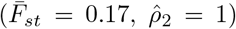 and rice 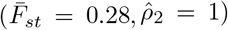. However, of apple 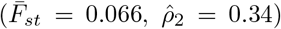 and soybean 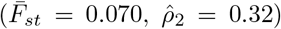, which had lower population structures, their respective correlation between 𝓁_*ij*_ and *v*_*ij*_ was also weakened. LD-eReg also showed a continuous distribution of *ρ*_2_ (**Figure 5I**), where populations with low 𝓁_*ij*_ often had a small *ρ*_2_. The dynamics of how *ρ*_2_ changed in response to *peeling* can be found in **Figure S17**. Notably, in species with extremely poor model fitting, such as sheep and dog, the red points in the regression results were divided into several clusters, similar to the distorted pattern seen on chromosome 6 in UKBA (**Figure 3N, O**).

### 5.2 Case studies for *RefPop*

Having reviewed the Norm I and Norm II patterns in ***RefPop***, we found that species such as Thale cress samples fit well in its Norm I 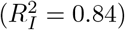 and Norm II (*ρ*_2_ = 0.95) patterns (**Figure S15U** and **Figure S16U**), but more puzzles remained for many species. However, a certain degree of distortion could be resolved, which led us to explore their underlying biological factors.

#### 5.2.1 Dog – extremely high LD blocks

Dog chromosomes 9, 19, and 26 showed a high deviation from the expected Norm I pattern, even after adjustment for its top 28 eigenvalues (**Figure 6A**). Then in its Norm II pattern, any 𝓁_*ij*_ involved with chromosome 8, 9, 19, or 26 was obviously separated from most of the chromosomal pairs (**Figure 6B**). We illustrated LD for chromosomes 8, 9, 19, and 26 (1, 000 × 1, 000 SNPs), and found a very tight LD block on each of the four chromosomes (chr 8: 73.0-74.3Mb, chr 9: 8.6-17.8Mb, chr 19: 20.0-20.3Mb, and chr 26: 25.2-27.1Mb) (**Figure 6E-H**). After these four blocks were removed, the Norm II pattern no longer showed two separate clusters (**Figure 6D**), which revealed that the significant deviation in Norm II pattern in the dog could be attributed to these four blocks. Although these blocks resembled the LD blocks of the MHC/HLA region in humans, the dog’s MHC region is actually located on chromosome 12 (53). Specifically, for chromosome 8, the high-LD region contains IGHV genes (54). For the other three chromosomes, we did not find annotations that could explain the existence of such LD patterns in dogs. However, there were large haplotypes in these corresponding regions in each of the four chromosomes (**Figure 6E-H**).

**Figure 6:**
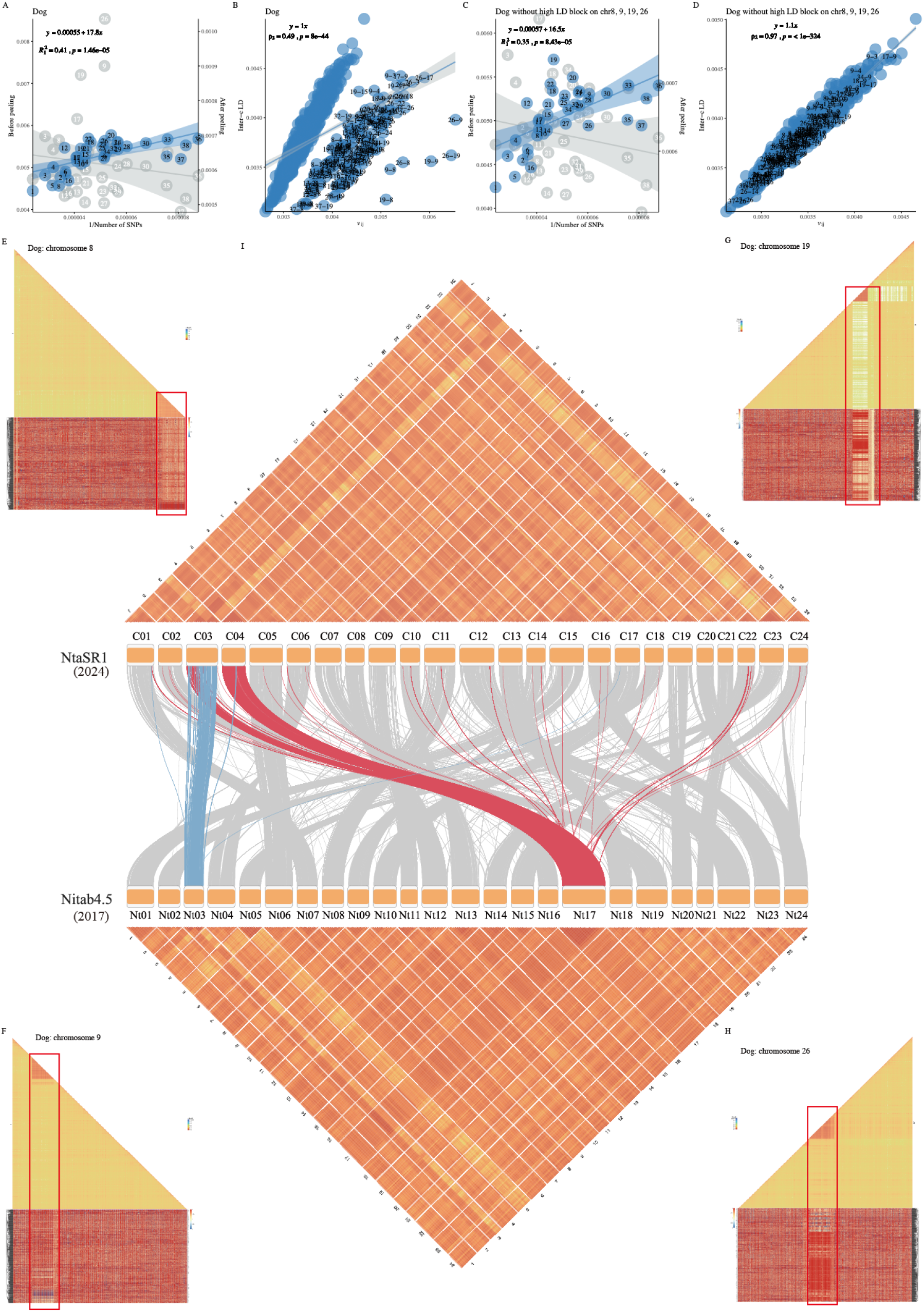
Investigation for the extremely high LD blocks in Dog and Tobacco. **A-B**, The Norm I and Norm II patterns in Dog. **C-D**, The Norm I and Norm II patterns in Dog after removing the extremely high LD blocks on chromosomes 8, 9, 19, and 26. **E-H**, High-resolution LD block illustrations for chromsome 8, 9, 19, and 26 in Dog with the corresponding haplotype plot beneath each illustration. **I**, High-resolution LD block illustrations for tobacco in NtaSR1 and Nitab4.5 and synteny analysis between them. The very large chromosome 17 in Nitab4.5 has largely been split into chromosomes 3 and 4 in NtaSR1.

In contrast, the sheep samples had a separated group in its Norm II pattern if chromosome 11 or 12 was involved (**Figure S16P**), but it was simply due to the very narrow genotyped SNPs in the short arms of chromosomes 11 and 12, respectively (**Figure S18**). Other obvious deviations were observed on chromosomes A06 and A08 in cotton (**Figure 5P**). Large genomic regions with lengths of about 40Mb on chromosome A06 and more than 70Mb on chromosome A08 shaped dramatic genomic divergence in cotton varieties planted in the Yangtze River region of China and northwest China. These large genomic regions with extremely LD have led to their unexpected high interchromosomal LD patterns with other cotton chromosomes (**Figure S20**).

#### 5.2.2 Tobacco – the puzzle of chromosome 17

In tobacco, there are now two publicly available reference genomes, Nitab4.5, which was released in 2017 with next-generation sequencing technology (52), and NtaSR1, which was released in 2024 with PacBio Hifi sequencing technology (37), respectively. There was a puzzle on chromosome 17 when mapping 13,339 tobacco SNPs to Nitab4.5, because not only chromosome 17 appeared to be unproportionally large (encompassing 16.4% of the total SNPs), and also chromosomes 3 and 17 exhibited a markedly high interchromosomal LD of 𝓁_3,17_ = 0.32, a substantial deviation in LD-eReg (**Figure S16V**). So, the extremely high value of 𝓁_3,17_ implied a plausibly incorrect alignment. After the release of NtaSR1, chromosome 17 in Nitab4.5 was predominantly aligned with chromosomes 3 and 4 in the NtaSR1 reference genome. Given the substantial colinearity between NtaSR1 and Nitab4.5, it was also found that part of chromosome 3 in Nitab4.5 was aligned into chromosome 3 in NtaSR1 (**Figure 6I**). Furthermore, the high-resolution LD block plot (100 × 100 SNP pairs for each block) was illustrated in perfect agreement with the mapping results; for example, the exceptionally high LD block on chromosomes 3 and 4 in NtaSR1 reflected with the corresponding mysteriously dense LD blocks on chromosome 17 in Nitab4.5.

## 6 Discussion

If LD analysis for a small region of a couple of hundreds of kilobase pairs has become routine, genomic LD that illustrates genome-wide LD pattern should be anticipated. The obstacle is obviously the substantial computational cost of *𝒪*(*n*^2^*m*) in its realization under conventional methods. We developed efficient X-LD/X-LDR algorithms that reduced the cost of computation and storage, the foundation was laid for the study of genomic LD in resequenced populations even with an extremely large sample size (**Figure 1**). We therefore embarked on an exploration for LD at an unprecedented scale in both sample size and species coverage. Most of the investigated datasets could have their high-resolution LD estimation completed in a couple of hours or even less, which should be considered affordable, and the most efficiency improvement could even be 10 ∼ 10^3^ times faster depending on the purpose of an investigation (**Table 1**). As a global inflator of LD (55), in this study the detailed mechanism by which the population structure shapes LD was derived, and *peeling* technique also was proved and demonstrated a useful approach in polishing the estimation of LD. These technical innovations will benefit many aspects of genetic studies in which LD is rooted.

Initially, we applied our efficient methods to human cohorts and found the neutrality of the human genome since its *de facto* standard Norm I pattern (*ρ*_1_ ≈ 1). The regression coefficient of LD-dReg reflected the well-known evolutionary timeline of African, European, and East Asian populations, respectively, and, furthermore, their analysis of LD decay also pointed to such an evolutionary timeline (**Figure 2**). Increasing rare variants obviously shaped LD patterns in UKB and led to stronger LD due to rare variants, consistent with recent investigation of much higher estimation of missing heritability by including more rare variants in TopMed (47). The distortion of Norm I was leveraged to pinpoint factors that distort LD patterns in human populations, and the main disturbance of the Norm I pattern appeared to be the population structure, which smeared the Norm I pattern but sharpened the Norm II pattern. Dividing the human genome into functional categories, the ncRNA regions exhibited exceptionally high LD in four independent human datasets, consistent with previous findings of such phenomenon (49). The pericentromeric regions, one of which led to upward deviation of chromosome 11 in the Norm I pattern, were not globally as strong as the population structure. The HLA region was another player, but it was not as strong as the pericentromeric regions. The selection of *FADS2* left footprint in at least Asian and European populations, but it did lift LD around *FADS2* but nearly ignorable compared to the centrometric region of chromosome 11. Intriguingly, polygenic selection for human height reduced the LD of the QTL regions, not statistically significant compared to the control regions, but could be expected from the Hill-Robertson effect (56; 57; 58). As T2T solved high-density alpha satellite DNA, which is composed of an AT-rich family of ∼171 bp monomers and repeated for even millions of bases (51), T2T reasonably strengthened the LD that *b*_1_ of LD-dReg increased from 53.2 to 83.3, a nearly 60% increase (**Figure 3**). In summary, the genomic LD patterns of the human population were influenced to some extent by factors of the population structure *>* pericentromeric region *>* HLA *>* polygenic selection, a hierarchy that was empirically found to be consistent in each investigated human ethnicity. However, the population structure could be efficiently removed using the peeling technique. Among the 22 human autosomes, chromosome 11 always stood out in both the pre-T2T and T2T genomes probably because of its large centrometric region. As expected, the behavior of human LD demonstrated human genome as an ideal thermodynamic gas given its Norm I pattern, and promised a step-by-step investigation when its Norm I was disturbed.

In contrast, ***RefPop*** was possibly characterized by nonrandom mating due to experimental manipulation, which significantly inflated LD. *Peeling* did help polish the Norm I pattern for ***RefPop***, but the increase was not as straight as in the 1KG human cohort compared. In ***RefPop*** the disturbing factors could be various. Using our algorithm, we efficiently detected epistatic interactions throughout the genome. For example, on chromosome 3 of maize (Figure 4**K**), we identified SNPs around 65Mb and 100Mb that exhibit extremely high LD (*R*^2^ ≥ 0.9). This unusually extended LD suggested the presence of co-evolving genes at these distant loci, which may be involved in shared biological pathways or functional interactions. For sheep population, the outstanding of chromosomes 11 and 12 both in the Norm I and Norm II patterns were due to its extremely few assayed SNPs on short arms only, an obvious technical issue that can be fixed later. For the cotton population, chromosomes A06 and A08 were remotely spotted in its Norm II pattern, and this phenomenon was possible due to strong selection trunks on these two chromosomes as previously reported (22). For the dog population, chromosomes 8, 9, 19, and 26 were found to have obviously strong LD blocks, which were never documented in dog GWAS studies (23). Considering genomic LD an omnibus statistic that manifests various biological events, a far-reaching application than demonstrated should be possible after our proposed algorithms make the previous infeasible LD computation affordable.

## 7 Materials and Methods

### 7.1 Decomposing genomic LD into intrachromosal and interchromosal components

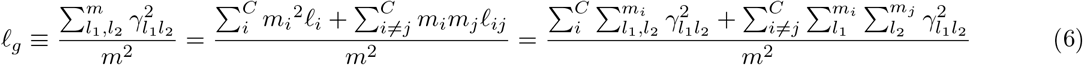

We define 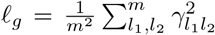 the genomic LD, which is the mean LD of *m*^2^ SNP pairs. Furthermore, we define the intrachromosomal LD component on a chromosome

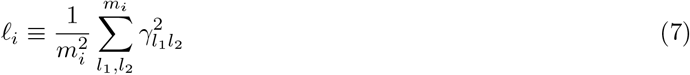

which averaged LD for the SNPs on the *i*^th^ chromosome, and define the interchromosomal component

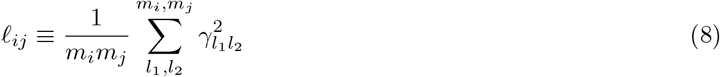

which averaged the pairwise LD for SNP *l*_*i*_ from the *i*^th^ chromosome and SNP *l*_*j*_ from the *j*^th^ chromosome. The decomposition is as schematically illustrated in **Figure 1**.

Using a conventional computational strategy such as that of PLINK, the computational cost is 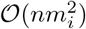 for **Equation 7**, or *𝒪*(*nm*_*i*_*m*_*j*_) for **Equation 8**. This computational cost is quickly infeasible for a resequencing population. To feasibly estimate the average LD for intrachromosomal and interchromosomal SNP pairs, we propose two novel approaches that fit GWAS or NGS data, in which *m* ≫ *n* but *n <* 10^4^ (X-LD) or *n* is extremely large (X-LDR), significantly alleviating both computational cost and storage. For more details on X-LD, please refer to Huang et al (9).

### 7.2 Method I: X-LD for small sample sizes

Given a standardized genotypic matrix **X**, a *n* × *m* matrix represents *n* individuals and *m* SNPs. Each SNP is scaled as 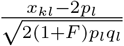, where *x*_*kl*_ is the genotype for the *k*^*th*^ individual at the *l*^th^ biallelic locus, *F* is the inbreeding coefficient having the value of 0 for the random mating population and 1 for the inbreed population, and *p*_*l*_ and *q*_*l*_ are the frequencies of the reference and alternative alleles, respectively. If we assume that there are *m*_*i*_ SNPs on chromosomes *i*, we can construct its corresponding *n* × *n* GRM,

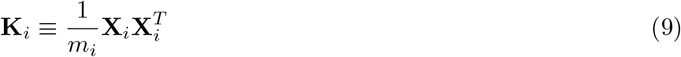

If we define the off-diagonal elements in Equation 9 as 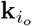 and 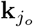, after some algebra, we have

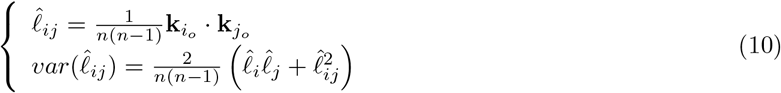

We can get the 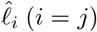 and 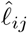, the intrachromosomal and interchromosomal LD, and their sampling variance. More details of X-LD can be found in (9).

### 7.3 Method II: X-LDR for biobank-scale sample

As the sample size increases, particularly as biobank-scale data become accessible, the approach to computing LD through Method I becomes infeasible. Here, a randomized estimator, denoted as *L*_*B*_, is applied, bypassing the cost of the construction of GRM (59). Specifically, for chromosomes *i* and *j*, we can define the following randomized estimators

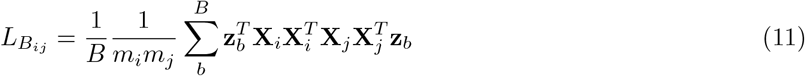

in which *B* is the round of iterations, **z**_*b*_ a vector of length *n* that is sampled from *N* (0, 1). As 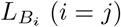 and 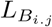 are the unbiased estimates of 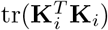, and 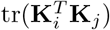, respectively. After further expanding 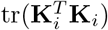, and 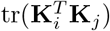 by Isserlis’ theorem (41), we can get 𝓁_*i*_ and 𝓁_*ij*_.

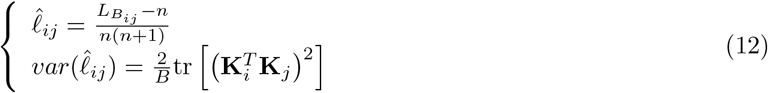

Besides, we can get the sampling variance of the estimated LD based on the intrachromosomal and interchromosomal LD in relation to 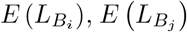, and 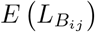. As the sampling variance of the estimated average LD is also related to sample size, it is critical to have a large sample size, a condition that distinguishes the application of Method I for small sample sizes and Method II for large sample sizes, such as UK Biobank.

Regardless of how the average LD is estimated, we can apply the two algorithms to estimate intra-*c* and inter-*c* LD patterns. For Method I, given *c* chromosomes, the computational cost is *𝒪*(*n*^2^*m*), in which 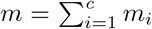 for Method II, the computational cost is *𝒪*(*nmB*). Using the Mailman algorithm can further improve the multiplication of the matrix in Equation 9 and Equation 11 (60). These optimization steps have been realized in our software.

### 7.4 Theoretical reconciliation for LD estimations

It is known that LD can be estimated in either phased, hereby represented as *γ*^2^, or unphased data, hereby *r*^2^, and their difference is reconciled that 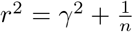, in which n is the sample size(9; 61). Otherwise specified, in this study the LD is referred to *γ*^2^ as estimated in unphased genotype data. In addition, caveats should be taken into account for the consistency between X-LD and X-LDR. Firstly, as either selection or diversity of samples would inflate *F*, in the step of standardization for genotypes, for some datasets, taking the inbreeding factor *F* would lead to inconsistency between XLD and XLDR. In this study, we always took *F* into account rather than considering it zero. Secondly, since X-LDR is a randomized algorithm, its estimates of LD exhibit higher variance, especially when sample sizes are small, compared to X-LD. Therefore, when sample sizes are not very large and X-LD can be estimated within a reasonable time, we recommend using X-LD.

### 7.5 The mechanism of *peeling*

For a pair of SNPs, with alleles 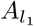 and 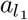 at locus *l*_1_, and 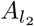 and 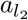 at locus *l*_2_, the association between the pair of SNP 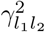 has a very general expression

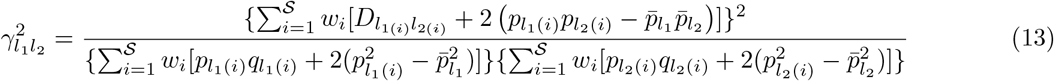

There is a “true” LD component 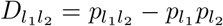, denoted as 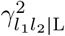, and can be written as

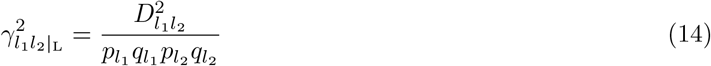

where *p*_*l*_. and *q*_*l*_. are the frequencies of the *l*^th^ reference and the alternative alleles in the population (62).

However, the population has 𝒮 subpopulations. If we let *n*_*i*_ be the size of the *i*^th^ subpopulation with 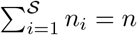 and 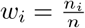 be the proportion of the *i*^th^ subpopulation. How to find these thresholds that divide individuals into 𝒮 subpopulations? Probably there are no such clear thresholds, but we can approximate them via recurrent dichotomization, the intuition of which lies in the established result 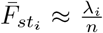 (43). 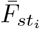 is the averaged 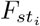 of all loci if the population is divided over the eigenvector that is associated with *λ*_*i*_ the *i*^th^ largest eigenvalue. Consequently, upon whether each individual’s coordinate is positive or negative on an eigenvector, the individuals can be divided into one of the two groups. So we can find a systematic way to dichotomize *n* individuals and eventually approximate the population structure. Now 𝒮 = 2, the allelic association due to population structure is

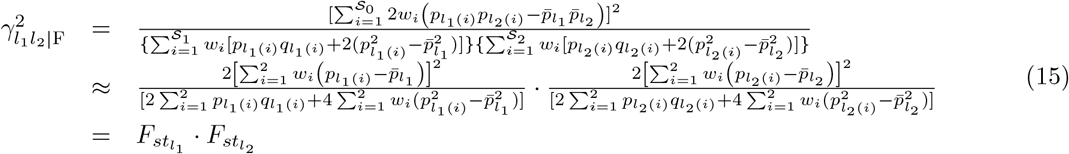

Now Equation 15 converts LD, the relationship between these two loci, into the production of their *F*_*st*_.

### 7.6 The number of eigenvalues that should be taken into account during peeling

We use *R*^2^ of LD-dReg to establish a robust *stopping rule* for determining the optimal number of eigenvalues to include. Specifically, starting from the first eigenvalue, we progressively incorporate eigenvalues into *peeling* model until reaching the point of maximum *R*^2^ or convergence. The number of eigenvalues at this stage is considered reasonable and denoted as *q*. Now, we are interested in 𝓁_*g*|F_ which averages all pairs of LD, and we have

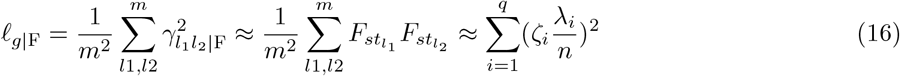

Of Equation 16, the first approximation takes through because 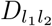 in Equation 13 only exists for the pair of SNPs nearby and their contribution can be ignored at the whole genome-wide scale. We argue that only the eigenvalue component greater than 1 of each selected eigenvalue should be utilized, which is a logical inference from EigenGWAS (63; 64; 65). In EigenGWAS, an eigenvector is regressed against each SNP, and the median of their corresponding 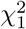 test has a value greater than 1 if there is population structure. Consequently, a weight 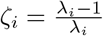 is derived to reduce the overcorrection.

For the calculation of eigenvalues, a full singular value decomposition (SVD) is used for datasets with a small sample size (*n <* 10^4^). For larger biobank-scale data, an Expectation-Maximization (EM) algorithm combined with the Mailman algorithm is utilized. The computational cost is expressed as 𝒪(*n*^2^*m*) and 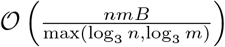 respectively, where *B* represents the number of iterations of the randomized algorithm (66).

## 8 Real datasets

A summary of the data used is presented in **Extended Data 1**. In particular, we highlight the data below for possible detailed reference.

### 8.1 Data description

#### 8.1.1 Westlake Biobank for Chinese

The Westlake Biobank for Chinese (WBBC) cohort which was launched to better understand the effect of genetic and environmental factors on growth and development from adolescents to adults (67). For now, it includes 4,480 participants (Han Chinese) that have undergone whole genome sequencing (WGS) at a mean depth of 14× using the NovaSeq 6000 system (Illumina Co.) (12; 15). For the raw dataset, only the SNPs located on the autosome and with missing genotype rates lower than 0.01 were retained. The SNPs were then categorized based on their MAF into common SNPs (MAF *>* 0.01) and rare SNPs (MAF ≤ 0.01). To dig deeper into the influence of rare SNPs on complex LD components, we meticulously stratified the SNPs into three distinct MAF bins (MAF ≤0.001, 0.001 *<* MAF *<*= 0.01, and 0.01 *<* MAF *<*= 0.5).

#### 8.1.2 CONVERGE for Chinese

CONVERGE (10,640 individuals), one of the largest whole genome sequencing cohorts, was used to investigate Major Depressive Disorder (MDD) in the Han Chinese population (13). To validate the role of rare SNPs on complex LD components in the Chinese population, we performed operations similar to WBBC for CONVERGE, consequently stratifying the SNPs into four MAF bins (MAF ≤ 0.0001, 0.0001 *<* MAF ≤0.001, 0.001 *<* MAF *<*= 0.01, and 0.01 *<* MAF *<*= 0.5).

#### 8.1.3 UK Biobank

Encompassing genotype data of approximately half a million individuals, UKB provides a novel reference panel to explore complex patterns of LD within large populations. In the dataset after imputation, we removed common variants with imputation score *<* 1 and rare variants with imputation score *<* 0.95. Then, for the entire dataset before and after genotype imputation, SNPs with missing genotype rates higher than 0.01 were eliminated, identical criteria were also employed for UKBB (UKB Field ID 22021 and 21000) and UKBW. Subsequently, for the datasets before genotype imputation, only SNPs common to the datasets after imputation were retained. Five MAF bins were set (MAF ≤ 0.00001, 0.00001 *<* MAF ≤ 0.0001, 0.0001 *<* MAF ≤ 0.001, 0.001 *<* MAF ≤ 0.01, and 0.01 *<* MAF ≤ 0.5) and three MAF bins (MAF ≤ 0.001, 0.001 *<* MAF ≤ 0.01, and 0.01 *<* MAF ≤ 0.5), respectively.

#### 8.1.4 Tobacco

A diverse panel of 164 tobacco accessions was selected from germplasm collections from the Guizhou Academy of Tobacco Science, Guiyang, Guizhou, China. Information on accessions, including names, collection sites, and categories, is provided in **Extended Data 2**. For each accession, we collected fresh young leaves and stored at -80^°^C until use. Genomic DNA was extracted for each sampled accession by a modified CTAB method. Then, a high-density tobacco 430K SNP array (Affymetrix, America) with SNP positions based on the Hongda reference genome was used for population genotyping. Due to chromosome loss, segmental deletion, frequent homoeologous recombination, the complex characteristics of allotetraploid, tobacco genomes tend to be unstable and difficult to assemble. To investigate the potential influence of these events on complex LD components, we mapped SNPs to the tobacco reference genome K326 according to the reference genome sequence Nitab4.5 around the SNP array (52) and NtaSR1 (37).

### 8.2 Quality control

We employed the same quality control metrics for datasets 1∼25. **i)** Only autosomal SNPs were retained; **ii)** SNPs with missing genotype rates greater than 0.2 were filtered; **iii)** Only SNPs with minor allele frequencies greater than 0.05 were retained.

### 8.3 Variation annotation

With regard to WBBC and CONVERGE, we first extracted SNPs that were shared by the two populations with MAF greater than 0.05, and the difference in MAF between the two populations was less than 0.1. Then, SNPs shared by WBBC, CONVERGE, and UKB white British were extracted, totaling 4,257,224 SNPs. To give a thorough understanding of the complex LD patterns at different functional segment scales, the SNP annotation was performed for WBBC, CONVERGE, and UKB white British populations according to the human genome GRCh37.p13 using the package ANNOVAR (48). Based on genome annotation, SNPs were mainly classified in downstream, upstream, exonic, intronic, ncRNA (exonic), ncRNA (intronic), UTR3, UTR5, and intergenic regions.

### 8.4 Data availability

Public genetic datasets used in this study can be freely downloaded from the following URLs.

**Apple**: https://ngdc.cncb.ac.cn/gvm/getProjectDetail?project=GVM000128

**Barley**: https://doi.org/10.5447/IPK/2018/9

**Cattle**: https://www.ebi.ac.uk/eva/?eva-study=PRJEB38336

**Chicken**: https://ngdc.cncb.ac.cn/chickensd/

**Common wheat**: https://wwwdev.ebi.ac.uk/eva/?eva-study=PRJEB31218

**Cotton**: http://120.78.174.209:30081/ftp

**Dog**: http://gong_lab.hzau.edu.cn/Animal_ImputeDB/#!/download_dog

**Fruit fly**: http://dgrp2.gnets.ncsu.edu/data.html

**Great tit**: https://datadryad.org/stash/dataset/doi:10.5061/dryad.p03j0

**Maize**: http://gong_lab.hzau.edu.cn/Plant_imputeDB/#!/download_maize

**Mouse**: https://www.ebi.ac.uk/eva/?eva-study=PRJEB45961

**Pig**: https://piggtex.farmgtex.org

**Rapeseed**: http://gong_lab.hzau.edu.cn/Plant_imputeDB/#!/download_brassica_napus

**Rice**: https://wwwdev.ebi.ac.uk/eva/?eva-study=PRJEB13618

**Sheep**: https://wwwdev.ebi.ac.uk/eva/?eva-study=PRJEB14685

**Silkworm**: https://db.cngb.org/search/project/CNP0002456/

**Sorghum**: https://www.ncbi.nlm.nih.gov/bioproject/?term=pRJNA612320

**Soybean**: http://bigd.big.ac.cn/gvm/getProjectDetail?project=GVM000063

**Tea**: https://data.mendeley.com/datasets/k627r5kj4d/3

**Thale cress 1000 Genomes Project**: https://wwwdev.ebi.ac.uk/eva/?eva-study=PRJNA273563

**Tobacco**: Extended Data 3 and 4 of this study.

**Tomato**: ftp://download.big.ac.cn/GVM/Solanum_lycopersicum/SNP/detailed_vcf/

**Water buffalo**: https://ngdc.cncb.ac.cn/gvm/getProjectDetail?project=GVM000043

**Yeast**: http://1002genomes.u-strasbg.fr/files/

**Human 1000 Genomes Project**: https://wwwdev.ebi.ac.uk/eva/?eva-study=PRJEB30460

**CONVERGE**: https://wwwdev.ebi.ac.uk/eva/?eva-study=PRJNA289433

**UK Biobank**: https://www.ukbiobank.ac.uk/

## Supporting information

LDAtlas_SuppFigs

ExtendData1

ExtendData2

GenotypeTobaccoEdwards

GenotypeTobacco2024

## 9 Software description

We have developed a versatile tool GEAR that packages X-LD and X-LDR algorithms to dissect complex LD patterns from chromosome-wise to regional levels. GEAR (here we refer to the complex LD pattern analysis) comprises three main modules: (1) Data management: we chose the PLINK compact binary file format (*.bed, *.bim, *.fam) as the input data format and provide basic quality control functions; (2) X-LD method which is suitable for small or median sample sizes; (3) X-LDR method which is suitable for biobank-scale sample sizes. For reducing memory consumption, a two-step method is recommended; The computer code of X-LD or X-LDR is written in C++ programming language and supports multithreading based on OpenMP for high-performance computing. More detailed information on the source code and documentation of GEAR can be found on GitHub (https://github.com/gc5k/gear2).

## 10 Acknowledgements

The authors thank the UK Biobank participants (UKB application 41376) and the Westlake Biobank participants for making this work possible. This work was partially supported by National Natural Science Foundation of China (31771392 to CGB, and 32061143019 and 82370887 to HFZ), CNTC (110202101032 (JY-09) to SY), and GZY-ZJ-KJ-23001 to GBC. Sponsors did not play a role in the design, preparation, and submission of the article. The authors thank Drs Chen Chengjie and Zhang Xintan for helpful discussion. The authors appreciate the support of high-performance computing from the Center for Bioinformatics and Big Data Technology at Zhejiang University and the High-performance Computing Center at Westlake University. The paper is dedicated to the memory of William G Hill for his pioneering work in linkage disequilibrium.

## 11 Author contributions

GBC conceived and initiated the study. TNZ, XH, and MYY conducted simulation and analyzed data; TNZ, XH, FL, QXZ, ZZ, WZ, and SY contributed to the curation of the data; GBC, TNZ, XH, QXZ, and GAQ developed the toolkit for the proposed methods; GBC and TNZ wrote the first draft of the article MYY and HFZ led the analysis in WBBC. All authors contributed to the writing, discussion of the article, and validation of the results.

## 12 Supplementary information

Supplementary Figure S1–19, Extended Data 1-4.

**Figure S1**: Reconciliation for linkage disequilibrium (LD) estimators in Yeast

**Figure S2**: Comparison for XLD vs XLDR in ***RefPop***

**Figure S3**: High-resolution LD block in UKBB-c and CONVERGE-c

**Figure S4**: Norm I patterns in the 26 cohorts of 1KG human

**Figure S5**: Norm II patterns in the 26 cohorts of 1KG human

**Figure S6**: EigenGWAS and high-resolution LD illustration for CONVERGE

**Figure S7**: The LD decay plot based on adjacent 100 LD blocks in CONVERGE-c, UKBB-c, and UKBW-c

**Figure S8**: *R*^2^ of the regression for matched blocks of two equal splits in CONVERGE-c, UKBB-c, and UKBW-c

**Figure S9**: The LD decay plot based on adjacent 100 LD blocks across CONVERGE-c, UKBB-c, and UKBW-c (based on new 1,783,915 common SNPs)

**Figure S10**: LD within and across MAF-bin for CONVERGE, WBBC, UKBW, and UKBB

**Figure S11**: The LD decay plot based on adjacent 100 LD blocks across five MAF bins in UKBW **Figure S12**: The LD decay plot based on adjacent 100 LD blocks with the increasing imputed SNPs **Figure S13**: The contrast between WBBC-T2T and CONVERGE

**Figure S14**: The dynamics of 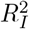 and *ρ*_1_ across ***RefPop***

**Figure S15**: LD atlas on the Norm I patterns in ***RefPop***

**Figure S16**: LD atlas on the Norm II patterns in ***RefPop***

**Figure S17**: The Norm I and Norm II dynamics in ***RefPop***

**Figure S18**: SNP density plot for sheep

**Figure S19**: Cotton haplotypes on A06 and A08

**Extended Data 1**: The results of LD-dReg across ***RefPop***

**Extended Data 2**: The sample ID and germplasm source of 164 tobacco accessions

**Extended Data 3**: The genotype data of 164 tobacco accessions after QC (Nitab4.5)

**Extended Data 4**: The genotype data of 164 tobacco accessions after QC (NtaSR1)

## References

[1] Slatkin, M. Linkage disequilibrium – Understanding the evolutionary past and mapping the medical future. Nature Reviews Genetics 9, 477–485 (2008).

[2] Miller, J. M., Poissant, J., Kijas, J. W., Coltman, D. W. & International Sheep Genomics Consortium. A genome-wide set of snps detects population substructure and long range linkage disequilibrium in wild sheep. Molecular Ecology Resources 11, 314–322 (2011).

[3] Rohlfs, R. V., Swanson, W. J. & Weir, B. S. Detecting coevolution through allelic association between physically unlinked loci. American Journal of Human Genetics 86, 674–685 (2010).

[4] Koch, E., Ristroph, M. & Kirkpatrick, M. Long range linkage disequilibrium across the human genome. PLoS ONE 8, e80754 (2013).

[5] Barrett, J. C., Fry, B., Maller, J. & Daly, M. J. Haploview: Analysis and visualization of LD and haplotype maps. Bioinformatics 21, 263–265 (2005).

[6] Dong, S. S. et al. LDBlockShow: A fast and convenient tool for visualizing linkage disequilibrium and haplotype blocks based on variant call format files. Briefings in Bioinformatics 22, 1–6 (2021).

[7] Chang, C. C. et al. Second-generation PLINK: Rising to the challenge of larger and richer datasets. GigaScience 4, 7 (2015).

[8] Theodoris, C., Low, T. M., Pavlidis, P. & Alachiotis, N. quickLD: An efficient software for linkage disequilibrium analyses. Molecular Ecology Resources 21, 2580–2587 (2021).

[9] Huang, X., Zhu, T.-N., Liu, Y.-C., Zhang, J.-N. & Chen, G.-B. Efficient estimation for large-scale linkage disequilibrium patterns of the human genome. eLife 12, 90636 (2023).

[10] Hoyt, S. J. et al. From telomere to telomere: The transcriptional and epigenetic state of human repeat elements. Science 376, eabk3112 (2022).

[11] The 1000 Genomes Project Consortium. A global reference for human genetic variation. Nature 526, 68–74 (2015).

[12] Cong, P. K. et al. Genomic analyses of 10,376 individuals in the Westlake BioBank for Chinese (WBBC) pilot project. Nature Communications 13, 2939 (2022).

[13] Cai, N. et al. Sparse whole-genome sequencing identifies two loci for major depressive disorder. Nature 523, 588–591 (2015).

[14] Bycroft, C. et al. The UK Biobank resource with deep phenotyping and genomic data. Nature 562, 203–209 (2018).

[15] Cong, P. et al. Identification of clinically actionable secondary genetic variants from whole-genome sequencing in a large-scale Chinese population. Clinical and Translational Medicine 12, e866 (2022).

[16] Lowy-Gallego, E. et al. Variant calling on the GRCH38 assembly with the data from phase three of the 1000 Genomes Project. Wellcome Open Research 4, 50 (2019).

[17] Liao, L. et al. Unraveling a genetic roadmap for improved taste in the domesticated apple. Molecular Plant 14, 1454–1471 (2021).

[18] Milner, S. G. et al. Genebank genomics highlights the diversity of a global barley collection. Nature Genetics 51, 319–326 (2019).

[19] Coppieters, W., Karim, L. & Georges, M. SNP-based quantitative deconvolution of biological mixtures: application to the detection of cows with subclinical mastitis by whole-genome sequencing of tank milk. Genome Research 30, 1201–1207 (2020).

[20] Wang, M.-S. et al. 863 genomes reveal the origin and domestication of chicken. Cell Research 30, 693–701 (2020).

[21] He, F. et al. Exome sequencing highlights the role of wild-relative introgression in shaping the adaptive landscape of the wheat genome. Nature Genetics 51, 896–904 (2019).

[22] He, S. et al. The genomic basis of geographic differentiation and fiber improvement in cultivated cotton. Nature Genetics 53, 916–924 (2021).

[23] Plassais, J. et al. Whole genome sequencing of canids reveals genomic regions under selection and variants influencing morphology. Nature Communications 10, 1489 (2019).

[24] Mackay, T. F. C. et al. The Drosophila melanogaster genetic reference panel. Nature 482, 173–178 (2012).

[25] Bosse, M. et al. Recent natural selection causes adaptive evolution of an avian polygenic trait. Science 358, 365–368 (2017).

[26] Bukowski, R. et al. Construction of the third-generation Zea mays haplotype map. GigaScience 7, gix134 (2018).

[27] Palma-Vera, S. E. et al. Genomic characterization of the world’s longest selection experiment in mouse reveals the complexity of polygenic traits. BMC Biology 20, 52 (2022).

[28] Teng, J. et al. A compendium of genetic regulatory effects across pig tissues. Nature Genetics 56, 112–123 (2024).

[29] Wu, D. et al. Whole-genome resequencing of a worldwide collection of rapeseed accessions reveals the genetic basis of ecotype divergence. Molecular Plant 12, 30–43 (2019).

[30] Wang, W. et al. Genomic variation in 3,010 diverse accessions of Asian cultivated rice. Nature 557, 43–9 (2018).

[31] Bolormaa, S. et al. Accuracy of imputation to whole-genome sequence in sheep. Genetics Selection and Evolution 51, 1 (2019).

[32] Tong, X. et al. High-resolution silkworm pan-genome provides genetic insights into artificial selection and ecological adaptation. Nature Communications 13, 5619 (2022).

[33] Lozano, R. et al. Comparative evolutionary genetics of deleterious load in sorghum and maize. Nature Plants 7, 17–24 (2021).

[34] Liu, Y. et al. Pan-genome of wild and cultivated soybeans. Cell 182, 162–176 (2020).

[35] Zhang, X. et al. Haplotype-resolved genome assembly provides insights into evolutionary history of the tea plant Camellia sinensis. Nature Genetics 53, 1250–1259 (2021).

[36] Alonso-Blanco, C. et al. 1,135 genomes reveal the global pattern of polymorphism in Arabidopsis thaliana. Cell 166, 481–491 (2016).

[37] Wang, J. et al. High-quality assembled and annotated genomes of Nicotiana tabacum and Nicotiana benthamiana reveal chromosome evolution and changes in defense arsenals. Molecular Plant 17, 423–437 (2024).

[38] Zhou, Y. et al. Graph pangenome captures missing heritability and empowers tomato breeding. Nature 606, 527–534 (2022).

[39] Luo, X. et al. Understanding divergent domestication traits from the whole-genome sequencing of swamp- and river-buffalo populations. National Science Review 7, 686–701 (2020).

[40] Peter, J. et al. Genome evolution across 1,011 Saccharomyces cerevisiae isolates. Nature 556, 339–44 (2018).

[41] Isserlis, L. On a formula for the product-moment coefficient of any order of a normal frequency distribution in any number of variables. Biometrika 12, 134–139 (1918).

[42] Zhou, Z.-Y. et al. PigVar: a database of pig variations and positive selection signatures. Database 2017, bax048 (2017).

[43] Patterson, N., Price, A. L. & Reich, D. Population structure and eigenanalysis. PLoS Genetics 2, e190 (2006).

[44] Ardlie, K. G., Kruglyak, L. & Seielstad, M. Patterns of linkage disequilibrium in the human genome. Nature Reviews Genetics 3, 299–309 (2002).

[45] Buckley, M. T. et al. Selection in Europeans on fatty acid desaturases associated with dietary changes. Molecular Biology and Evolution 34, 1307–1318 (2017).

[46] Liu, S. et al. Genomic analyses from non-invasive prenatal testing reveal genetic associations, patterns of viral infections, and Chinese population history. Cell 175, 347–359 (2018).

[47] Wainschtein, P. et al. Assessing the contribution of rare variants to complex trait heritability from wholegenome sequence data. Nature Genetics 54, 263–273 (2022).

[48] Wang, K., Li, M. & Hakonarson, H. ANNOVAR: functional annotation of genetic variants from highthroughput sequencing data. Nucleic Acids Research 38, e164 (2010).

[49] Vergara-Lope, A., Ennis, S., Vorechovsky, I., Pengelly, R. J. & Collins, A. Heterogeneity in the extent of linkage disequilibrium among exonic, intronic, non-coding RNA and intergenic chromosome regions. European Journal of Human Genetics 27, 1436–1444 (2019).

[50] Yengo, L. A saturated map of common genetic variants associated with human height. Nature 610, 704–12 (2022).

[51] Altemose, N. et al. Complete genomic and epigenetic maps of human centromeres. Science 376, eabl4178 (2022).

[52] Edwards, K. D. et al. A reference genome for Nicotiana tabacum enables map-based cloning of homeologous loci implicated in nitrogen utilization efficiency. BMC Genomics 18, 448 (2017).

[53] Miyamae, J. et al. Haplotype structures and polymorphisms of dog leukocyte antigen (DLA) class I loci shaped by intralocus and interlocus recombination events. Immunogenetics 74, 245–259 (2022).

[54] Rout, E. D., Burnett, R. C., Labadie, J. D., Yoshimoto, J. A. & Avery, A. C. Preferential use of unmutated immunoglobulin heavy variable region genes in Boxer dogs with chronic lymphocytic leukemia. PLoS ONE 13, e0191205 (2018).

[55] Nei, M. & hsiung Li, W. Linkage disequilibrium in subdivided populations. Genetics 75, 213–219 (1973).

[56] Hill, W. G. & Robertson, A. The effect of linkage on limits to artificial selection. Genetical Research 8, 269–294 (1966).

[57] Comeron, J. M., Williford, A. & Kliman, R. M. The Hill-Robertson effect: Evolutionary consequences of weak selection and linkage in finite populations. Heredity 100, 19–31 (2008).

[58] Charlesworth, B. et al. From Mendel to quantitative genetics in the genome era: the scientific legacy of W. G. Hill. Nature Genetics 54, 934–939 (2022).

[59] Wu, Y. & Sankararaman, S. A scalable estimator of SNP heritability for biobank-scale data. Bioinformatics 34, i187–i194 (2018).

[60] Liberty, E. & Zucker, S. W. The Mailman algorithm: A note on matrix-vector multiplication. Information Processing Letters 109, 179–182 (2009).

[61] Weir, B. S. & Hill, W. G. Effect of mating structure on variation in linkage disequilibrium. Genetics 95, 477–488 (1980).

[62] Weir, B. S. Linkage disequilibrium and association mapping. Annual Review of Genomics and Human Genetics 9, 129–142 (2008).

[63] Chen, G.-B., Lee, S. H., Zhu, Z.-X., Benyamin, B. & Robinson, M. R. EigenGWAS: finding loci under selection through genome-wide association studies of eigenvectors in structured populations. Heredity 117, 51–61 (2016).

[64] Galinsky, K. J. et al. Fast principal-component analysis reveals convergent evolution of ADH1B in Europe and East Asia. American Journal of Human Genetics 98, 456–472 (2016).

[65] Qi, G.-A. et al. EigenGWAS: An online visualizing and interactive application for detecting genomic signatures of natural selection. Molecular Ecology Resources 21, 1732–1744 (2021).

[66] Agrawal, A., Chiu, A. M., Le, M., Halperin, E. & Sankararaman, S. Scalable probabilistic PCA for large-scale genetic variation data. PLoS Genetics 16, 729202 (2020).

[67] Zhu, X. W. et al. Cohort profile: The Westlake BioBank for Chinese (WBBC) pilot project. BMJ Open 11, e045564 (2021).

