## Supplementary material for "An Atlas of Linkage Disequilibrium Across Species": LDAtlas_SuppFigs

September 24, 2024

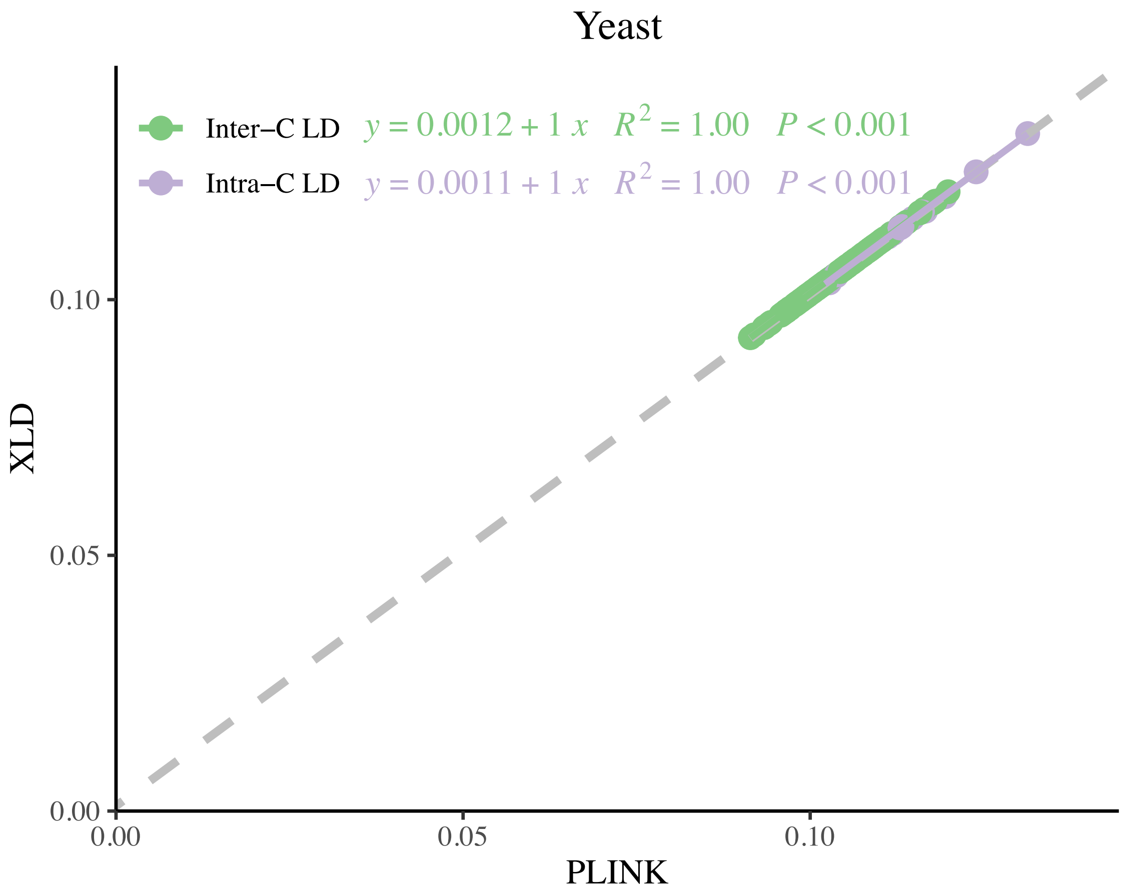


Figure S1: **Reconciliation for linkage disequilibrium (LD) estimators in Yeast** Consistency examination for yeast cohort for its $\hat{\ell}_i$ and $\hat{\ell}_{ij}$ estimated by X-LD and PLINK (–r2). The fitting line of $\hat{\ell}_i$ is purple, whereas the fitting line of $\hat{\ell}_{ij}$ is green. The gray dashed line, $y = \frac{1}{n} + x$, in which $n = 1,011$ the sample size of the yeast cohort. The two estimated regression models are at the top.

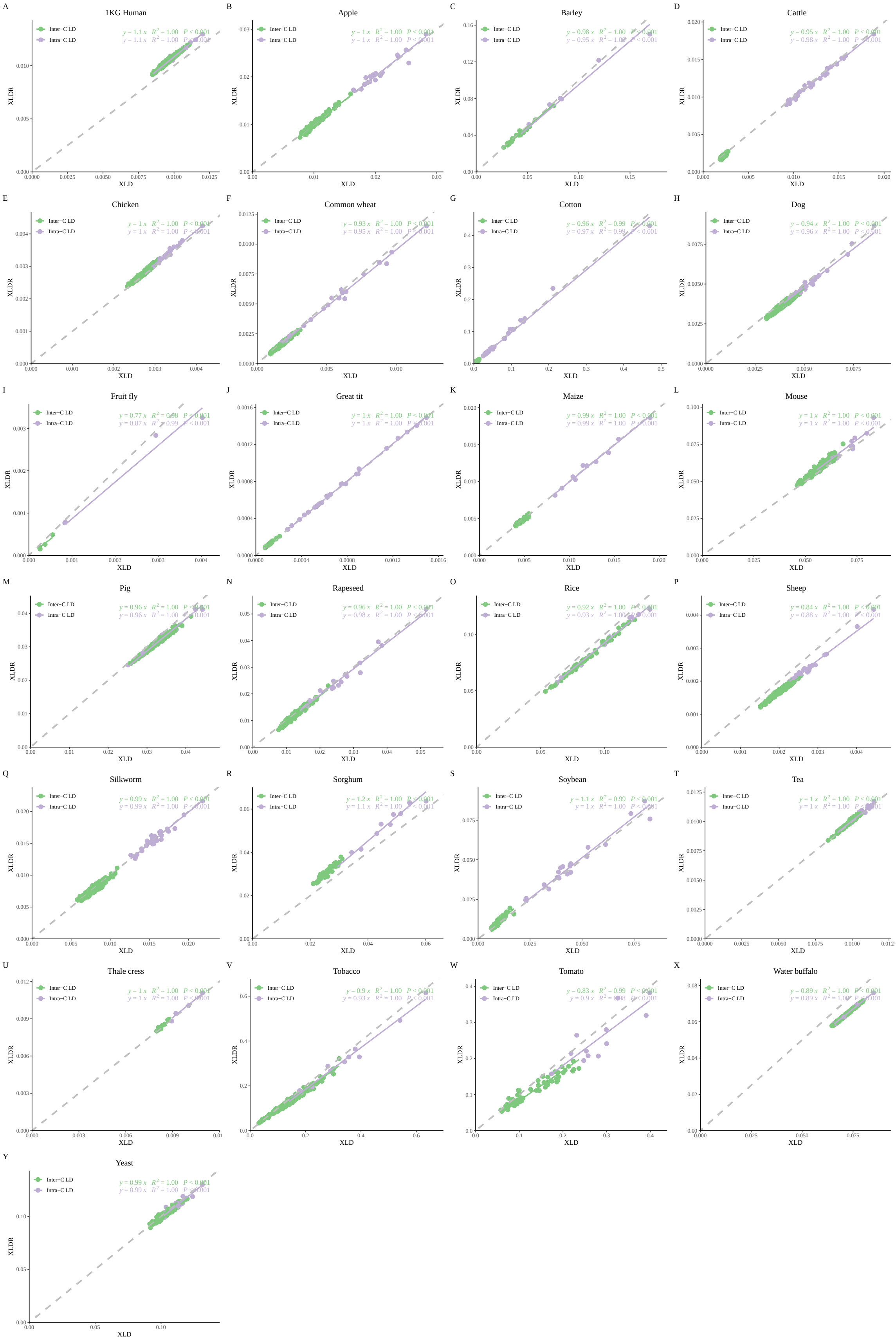


Figure S2: **Comparison for XLD vs XLDR in *RefPop*. A-Y**, Consistency examination for ***RefPop*** for their $\ell_i$ (intrachromosomal LD) and $\ell_{ij}$ (interchromosomal LD) are estimated by XLD (x-axis) and XLDR (y-axis, B=200 for XLDR). The similarity of the estimates between XLD and XLDR is represented by the linear regressions at the top-right corner of each plot. The fitting line of $\hat{\ell}_i$ is purple, whereas the fitting line of $\hat{\ell}_{ij}$ is green. The dashed diagonal line ($y = x$) is for reference.

A

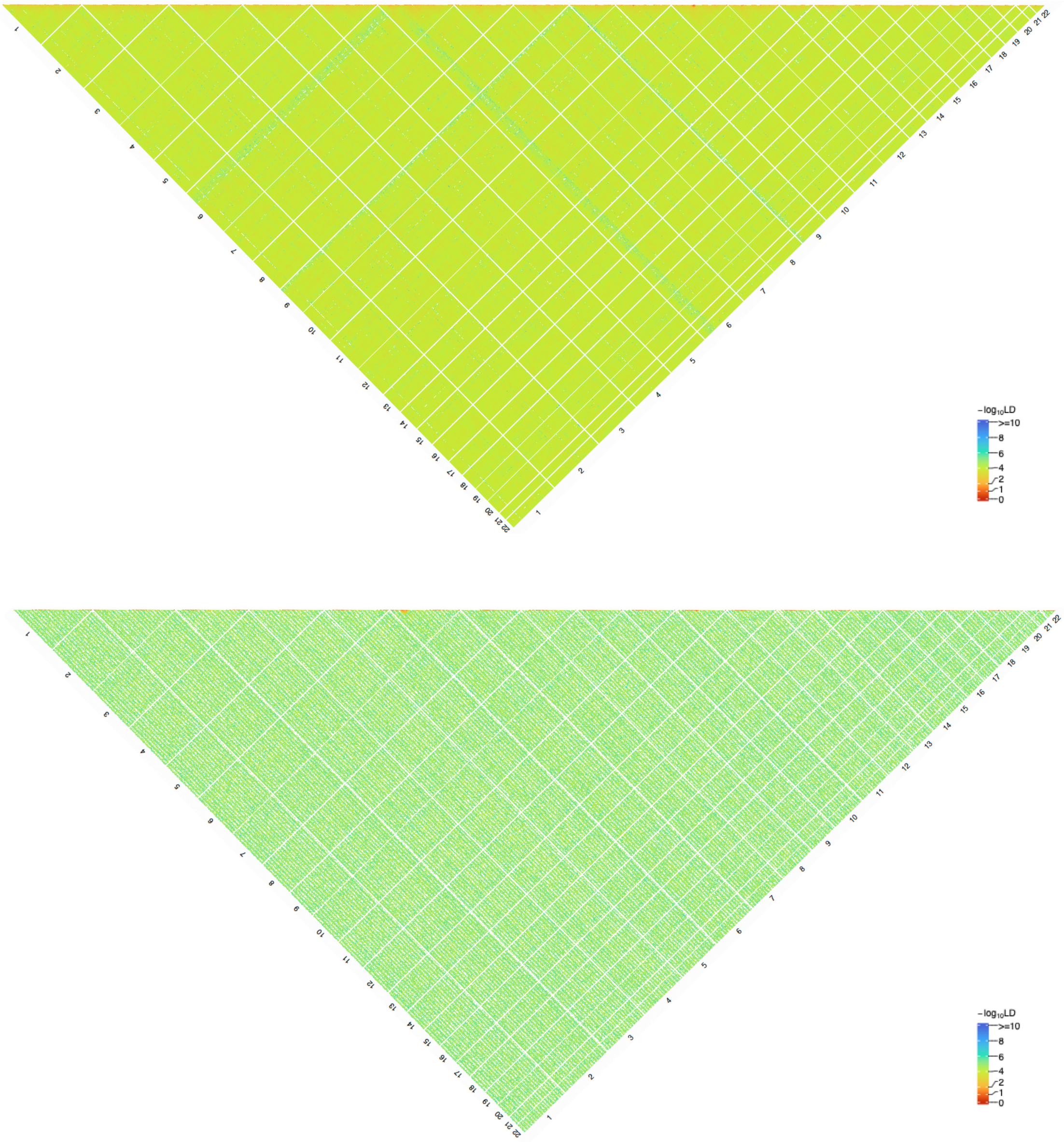

B

Figure S3: **High-resolution LD block in UKBB-c and CONVERGE-c**.
**A**, High-resolution LD illustration for UKBB-c. **B**, High-resolution LD illustration for CONVERGE-c.

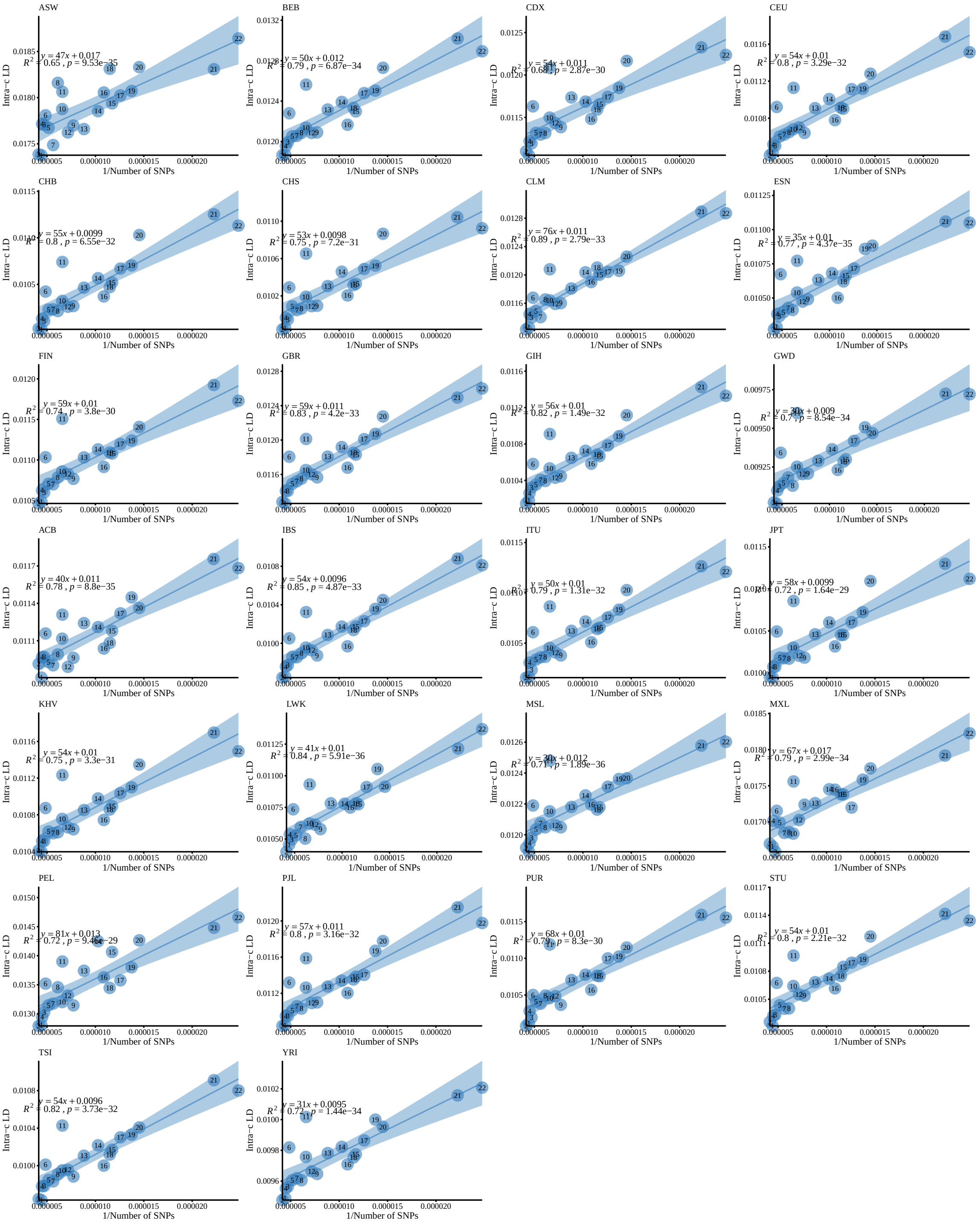


Figure S4: **Norm I patterns in the 26 cohorts of 1KG human.** The LD-dReg of each 1KG cohort without peeling.

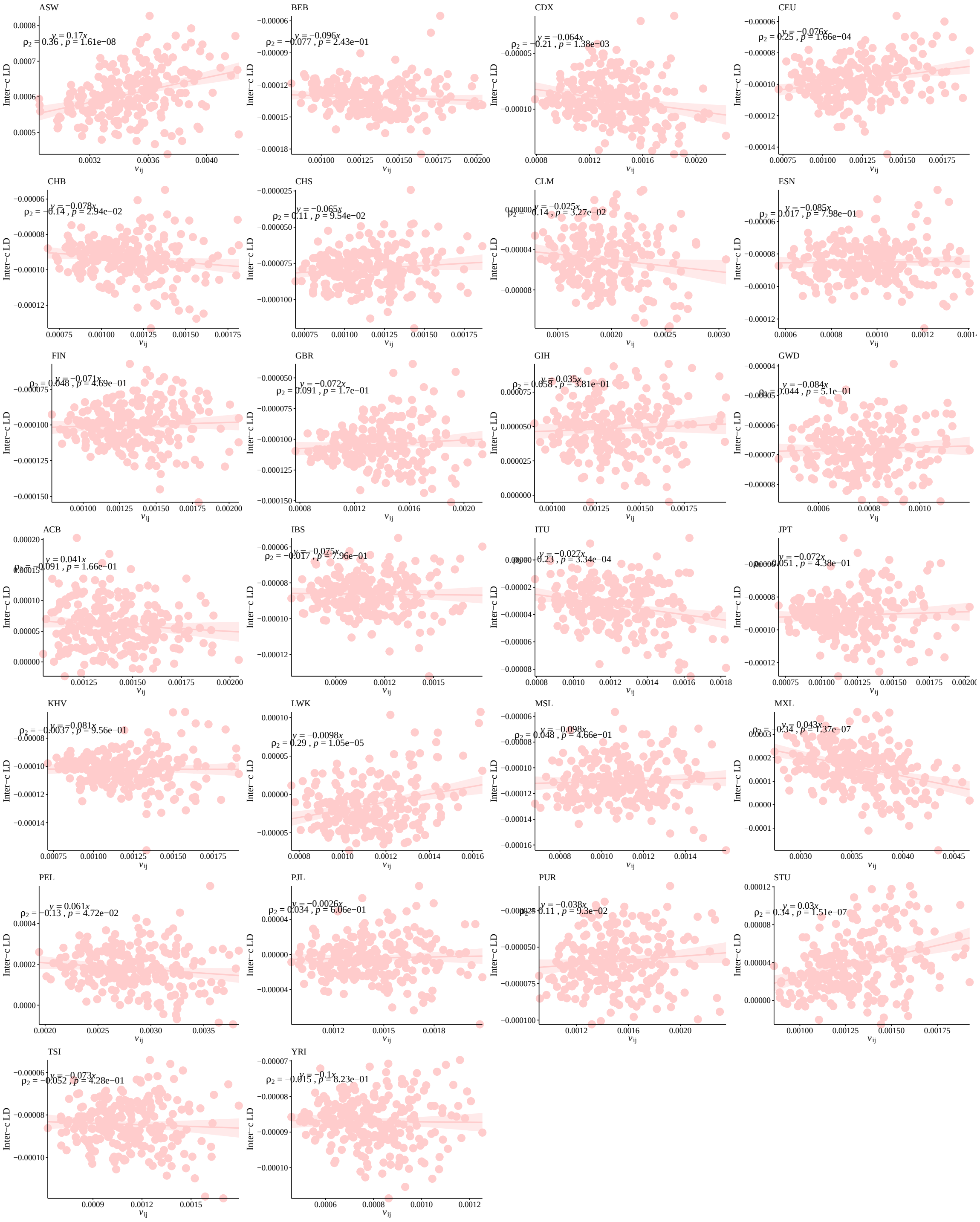

Figure S5: **Norm II patterns in the 26 cohorts of 1KG human.** The LD-eReg of each 1KG cohort.

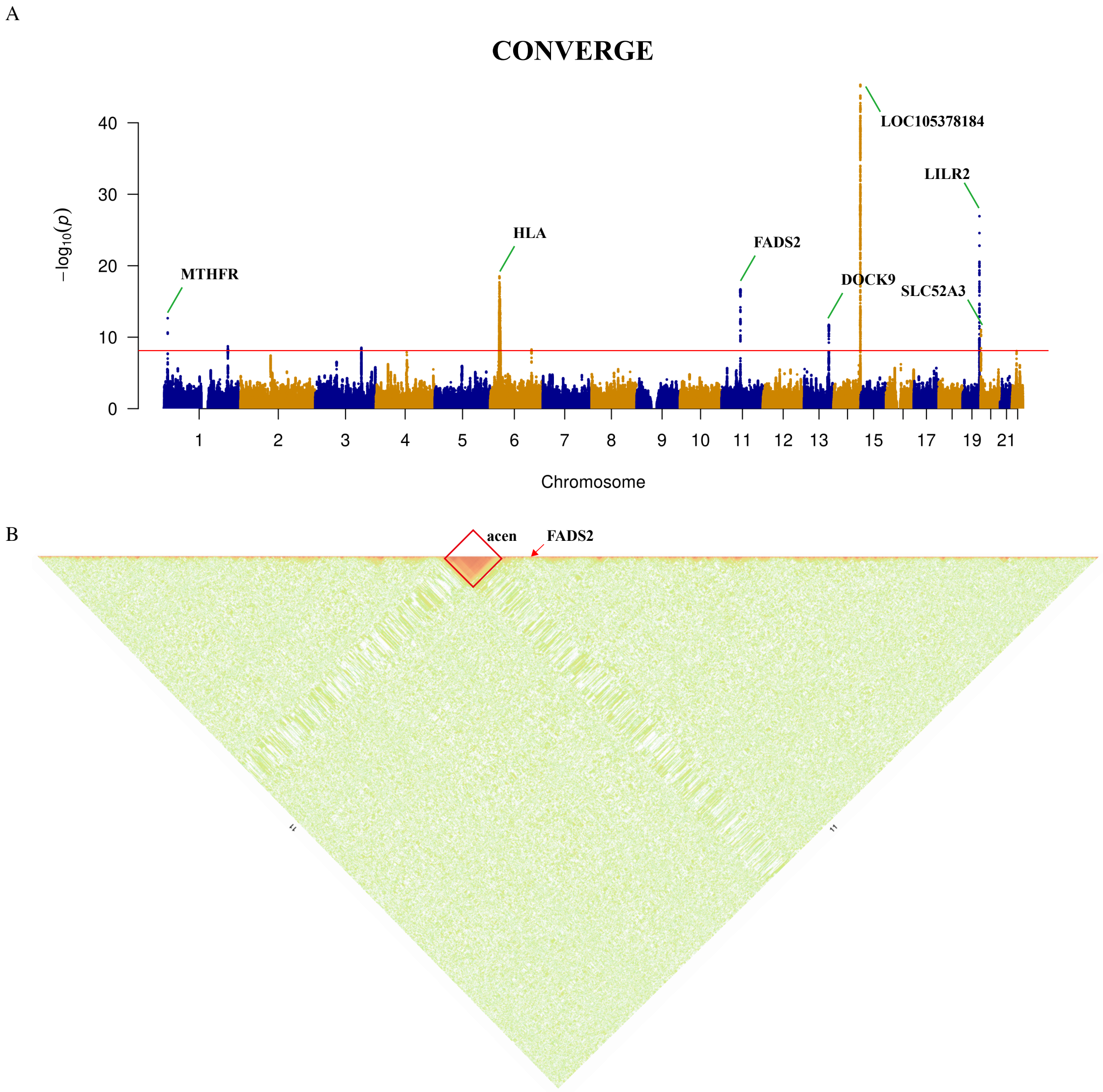


Figure S6: **EigenGWAS and high-resolution LD illustration for CONVERGE A**, the manhattan plot of EigenGWAS on the first eigenvector of CONVERGE. **B**, the high-resolution LD illustration for CONVERGE, the pericentromeric region and FADS2 region are annotated by red square and arrow, respectively.

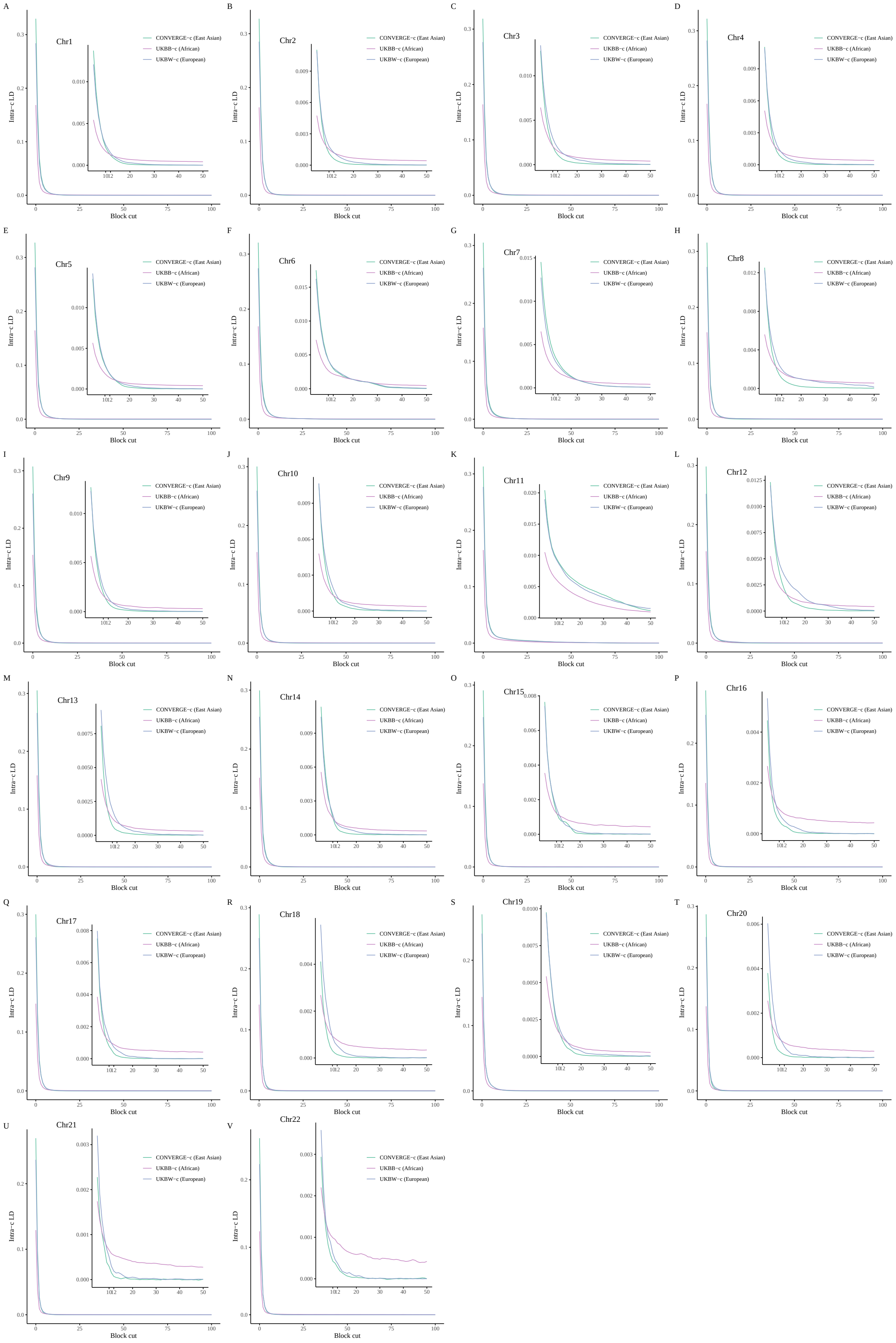


Figure S7: **The LD decay plot based on adjacent 100 LD blocks across CONVERGE-c, UKBB-c, and UKBW-c. A-V**, The LD decay plot from chromosome 1 to 22.

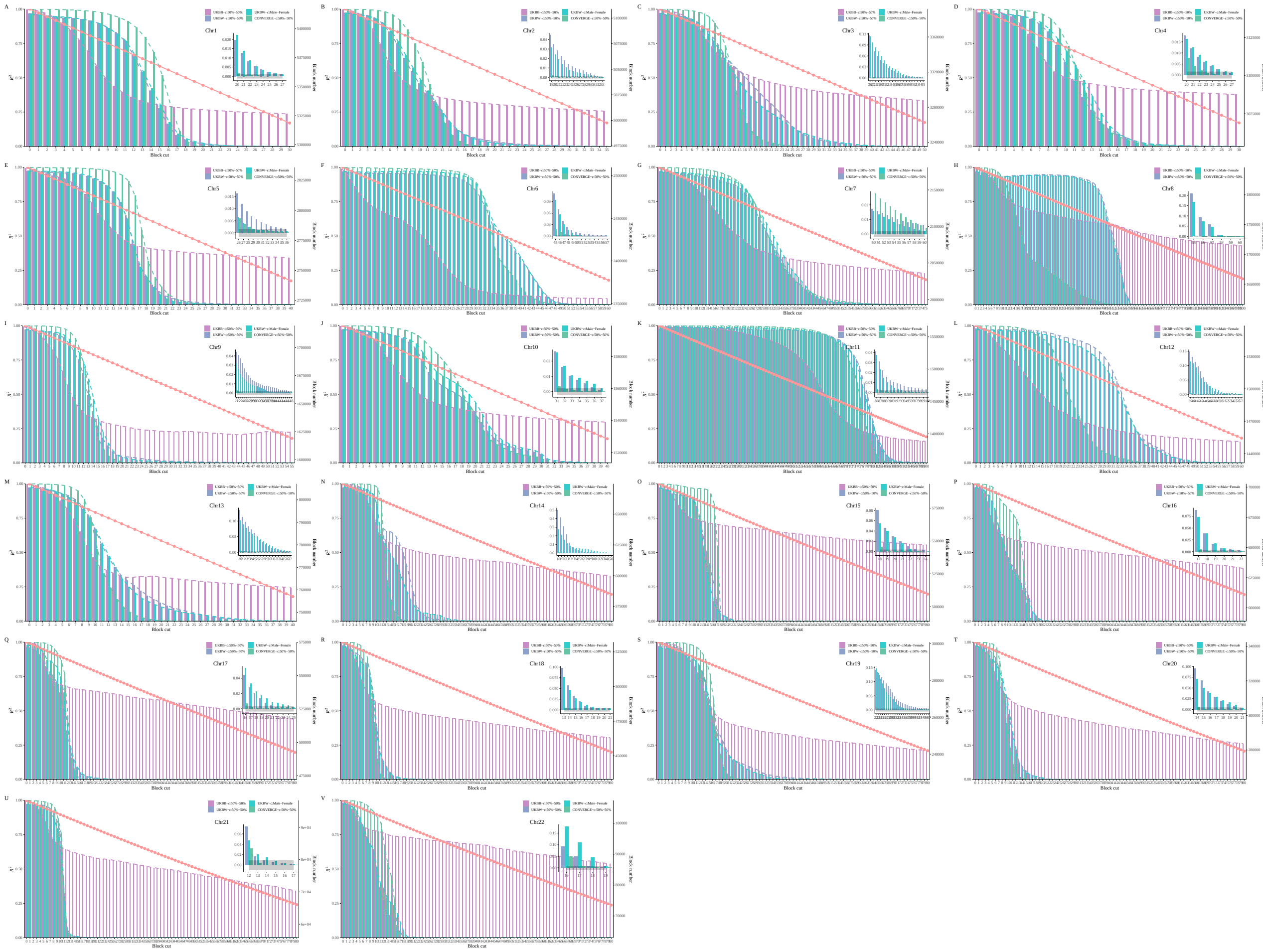

Figure S8: $R^2$ **of the regression for matched blocks of two equal splits in CONVERGE-c, UKBB-c, and UKBW-c A-Y**, $R^2$ of the regression for matched blocks of two equal splits across three single-ethnicity from chromosome 1 to 22; the pink points represent the number of blocks uesd in the regression.

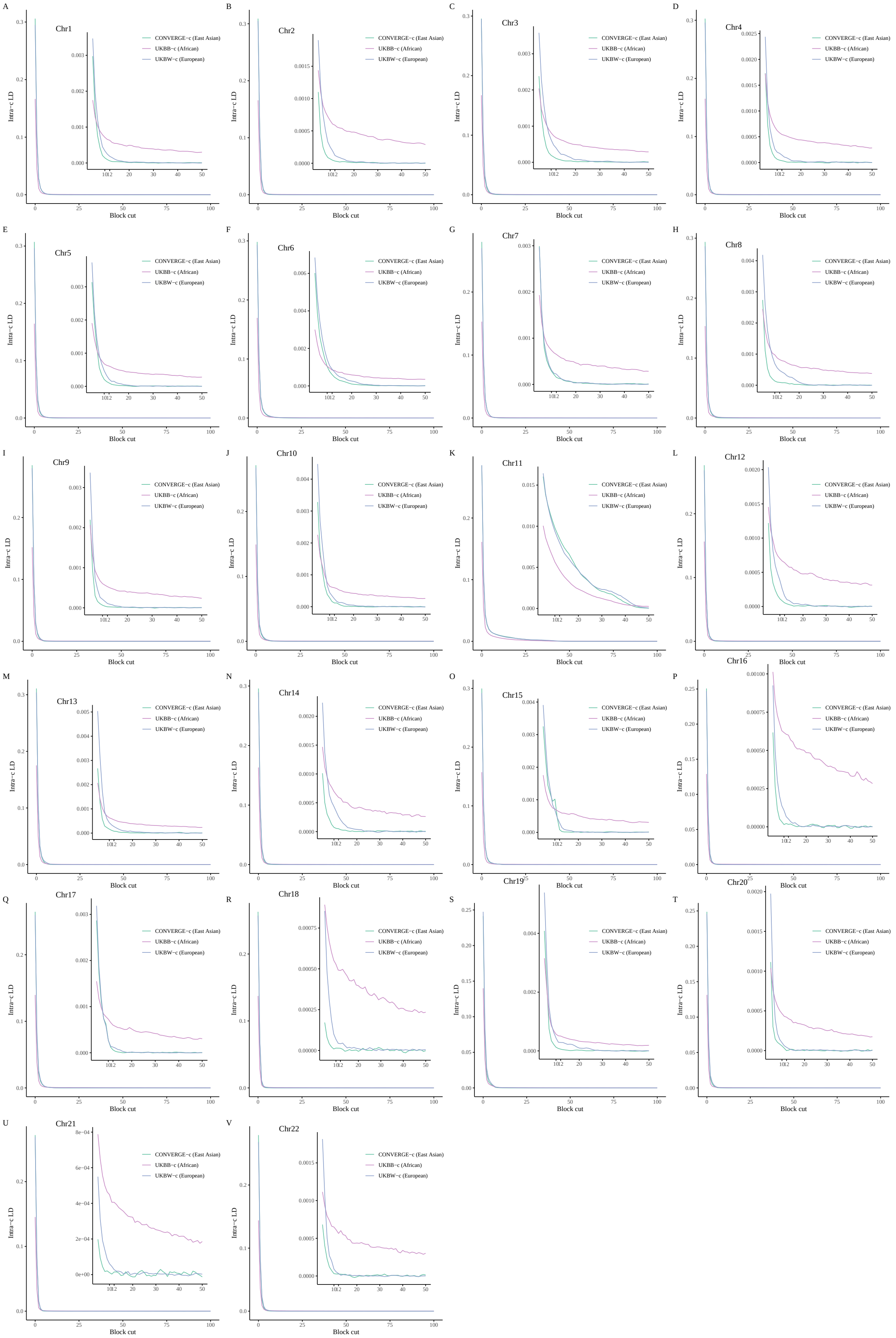


Figure S9: **The LD decay plot based on adjacent 100 LD blocks in CONVERGE-c, UKBB-c, and UKBW-c (based on new 1,783,915 common SNPs).A-V**, The LD decay plot from chromosome 1 to 22.

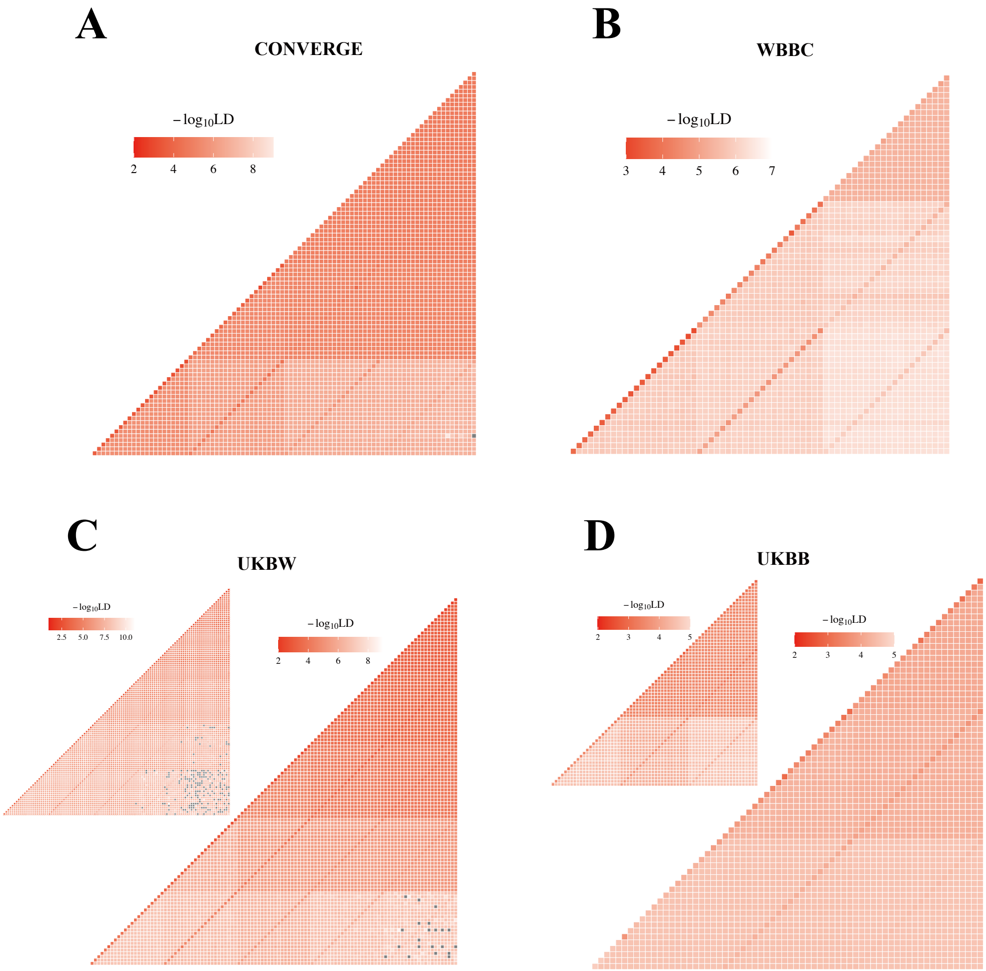


Figure S10: **LD within and across MAF-bin for CONVERGE, WBBC, UKBW, and UKBB. A**, four MAF bins were set from left to right ($0.01 < MAF \leq 0.5$, $0.001 < MAF \leq 0.01$, $0.0001 < MAF \leq 0.001$, $MAF \leq 0.0001$) for CONVERGE. **B**, three MAF bins were set from left to right ($0.01 < MAF \leq 0.5$, $0.001 < MAF \leq 0.01$, $MAF \leq 0.001$) for WBBC. **C**, five MAF bins were set from left to right ($0.01 < MAF \leq 0.5$, $0.001 < MAF \leq 0.01$, $0.0001 < MAF \leq 0.001$, $0.00001 < MAF \leq 0.0001$, $MAF \leq 0.00001$ ) for UKBW. (the result of UKBW chip dataset is on the topleft corner) **D**, three MAF bins were set from left to right ($0.01 < MAF \leq 0.5$, $0.001 < MAF \leq 0.01$, $MAF \leq 0.001$) for UKBB. (the result of UKBB chip dataset is on the topleft corner)

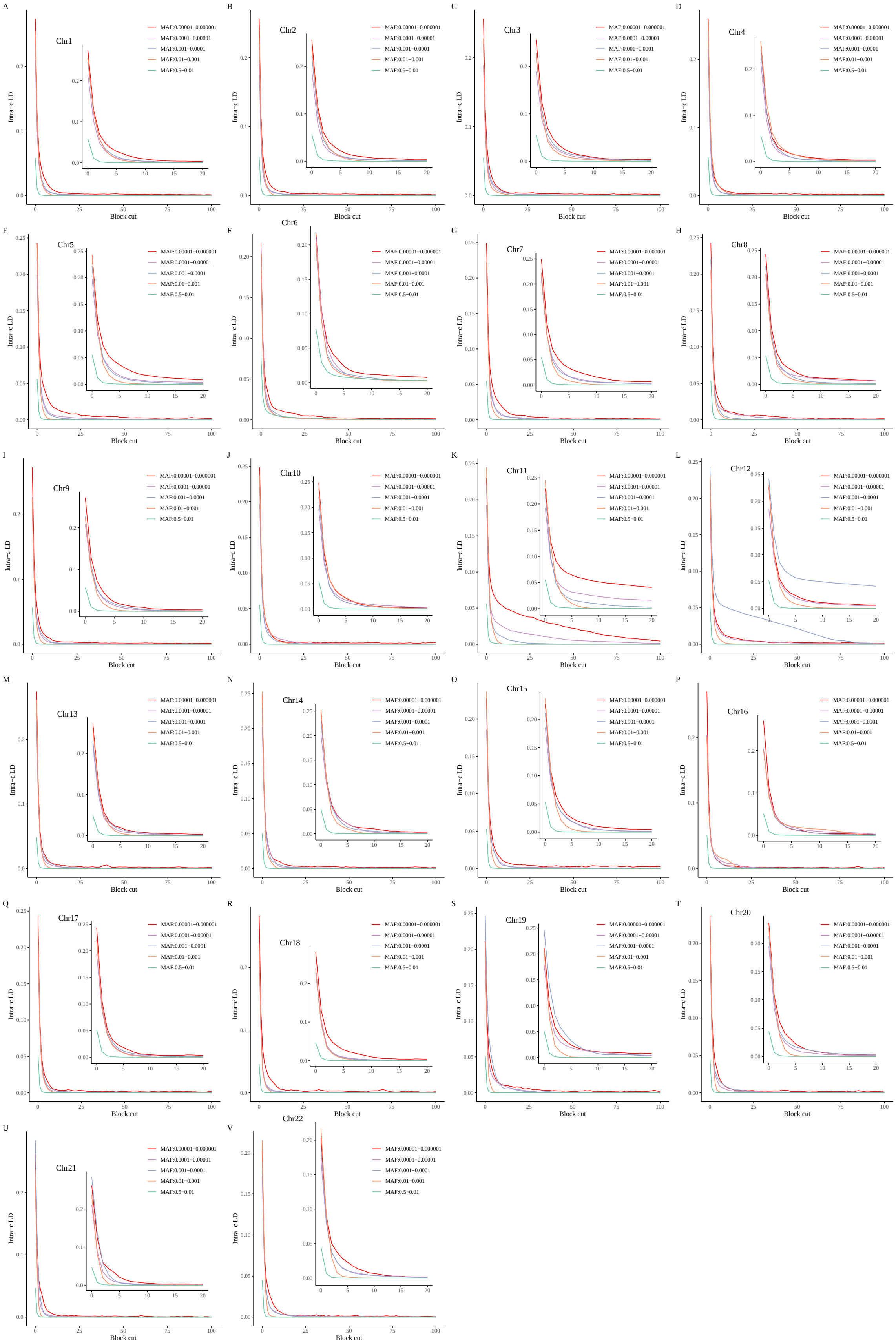


Figure S11: **The LD decay plot based on adjacent 100 LD blocks across five MAF bins in UKBW. A-V**: The LD decay plot from chromosome 1 to chromosome 22.

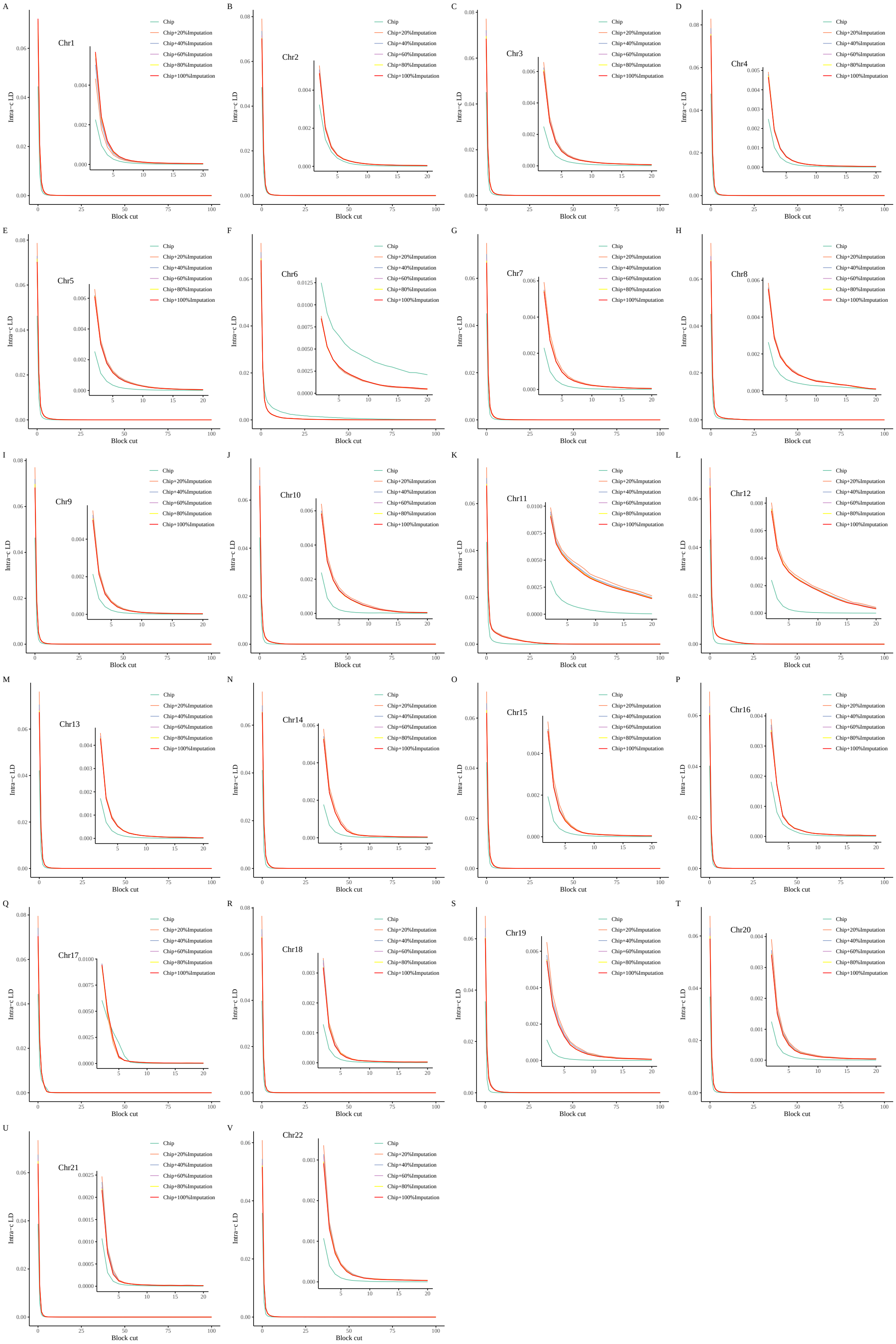

Figure S12: **The LD decay plot based on adjacent 100 LD blocks with the increasing imputed SNPs. A-V**: The LD decay plot from chromosome 1 to chromosome 22.

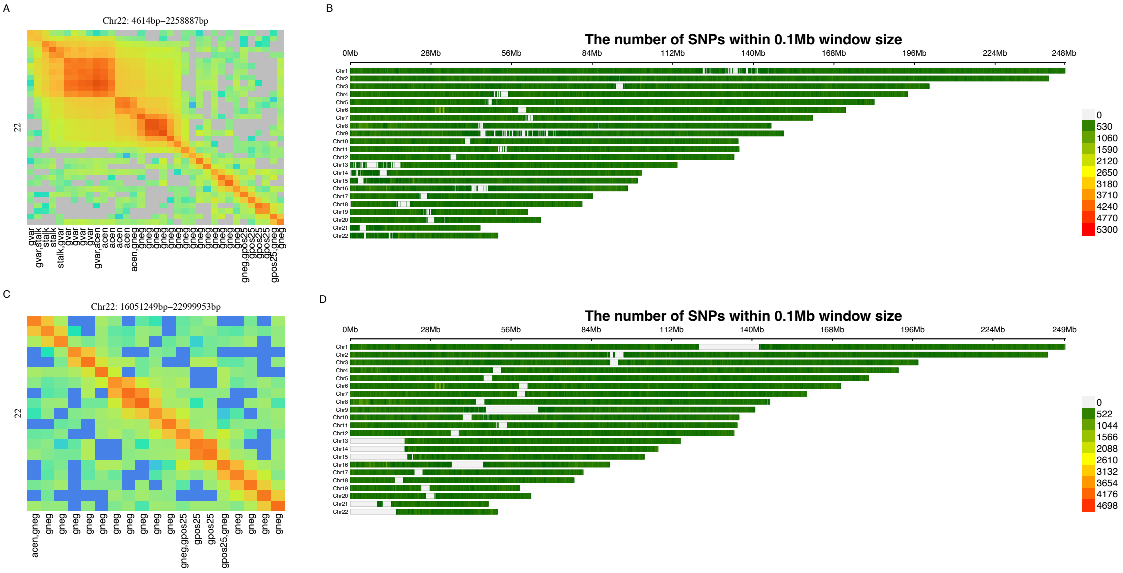


Figure S13: **The contrast between WBBC-T2T and CONVERGE.** **A**, High-resolution $\ell_i$ around the centromere region of chromsome 22 for WBBC-T2T (4,614-2,258,887bp). **B**, the SNP density plot for WBBC-T2T. **C**, High-resolution $\ell_i$ around the centromere region of chromsome 22 for CONVERGE (16,051,249-22,999,953bp). **D**, the SNP density plot for CONVERGE.

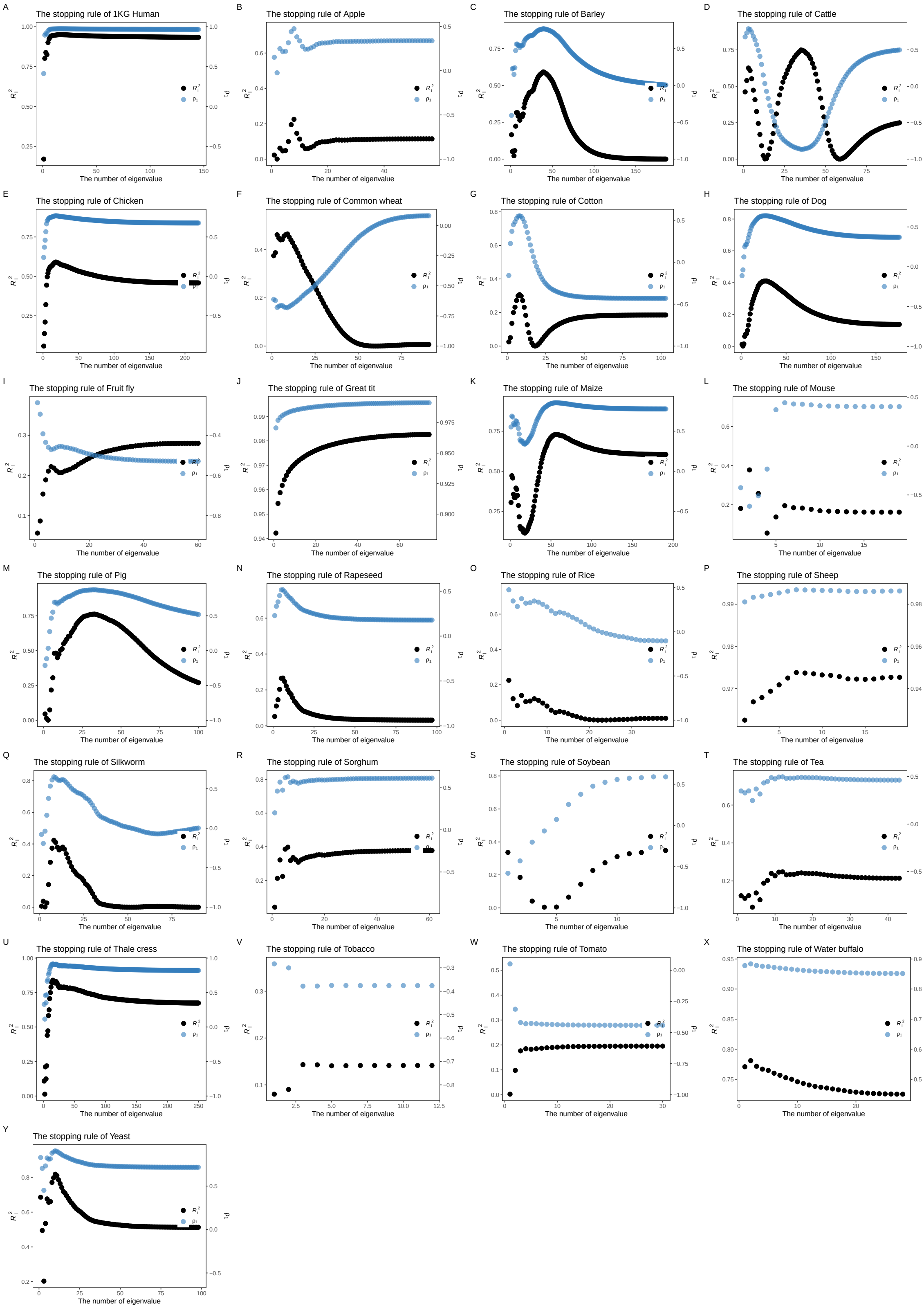


Figure S14: **The dynamics of $R_I^2$ and $\rho_1$ across *RefPop*. A-Y**, illustrations of the stopping rule across RefPop, which the black and blue points represent the $R_I^2$ and the $\rho_1$ of the Norm I pattern, respectively.

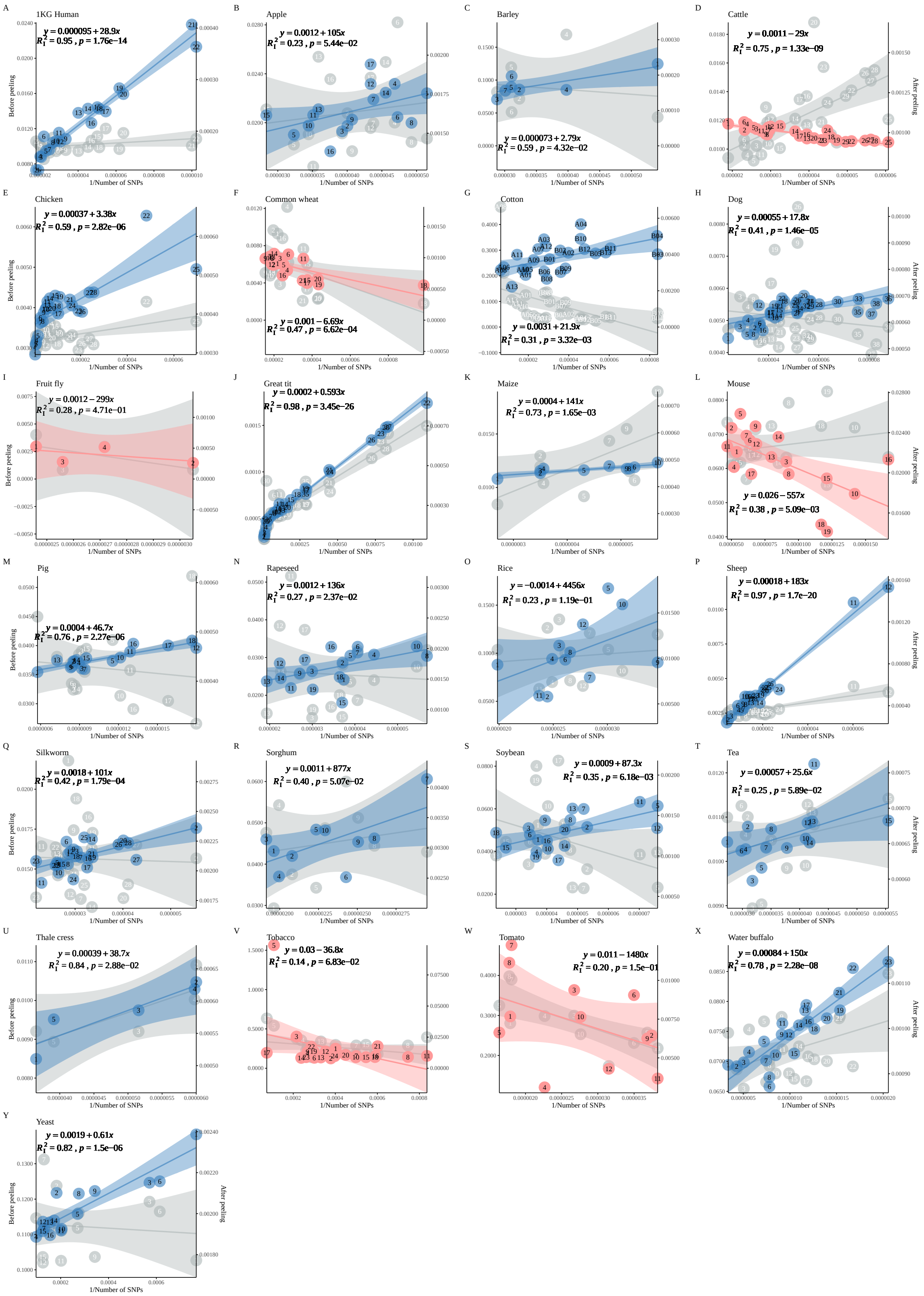

Figure S15: **LD atlas on the Norm I patterns in *RefPop*. A-Y**, The LD-dReg before and after *peeling* in ***RefPop***.

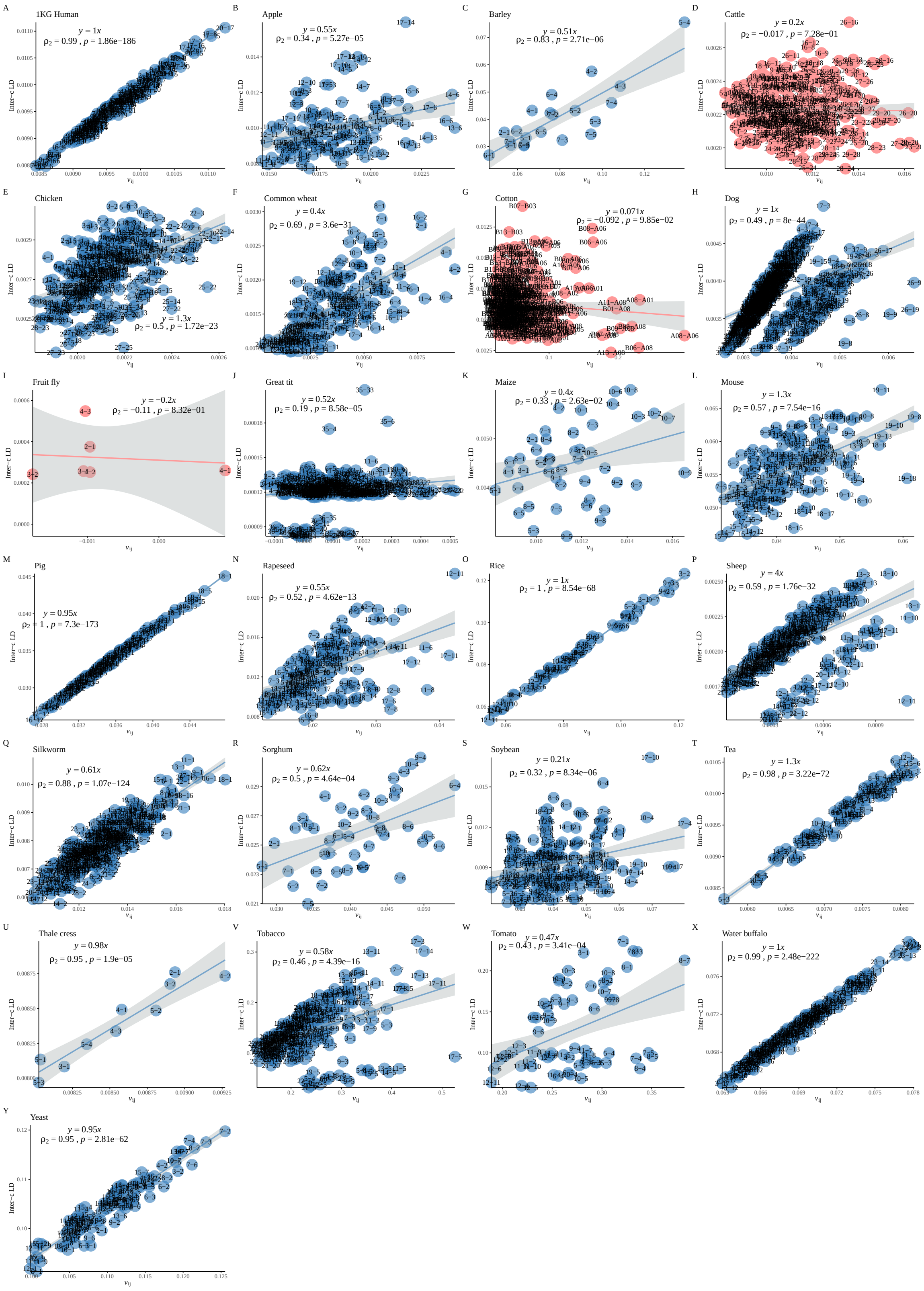


Figure S16: **LD atlas on the Norm II patterns in *RefPop*. A-Y**, The LD-eReg before and after *peeling* in ***RefPop***.

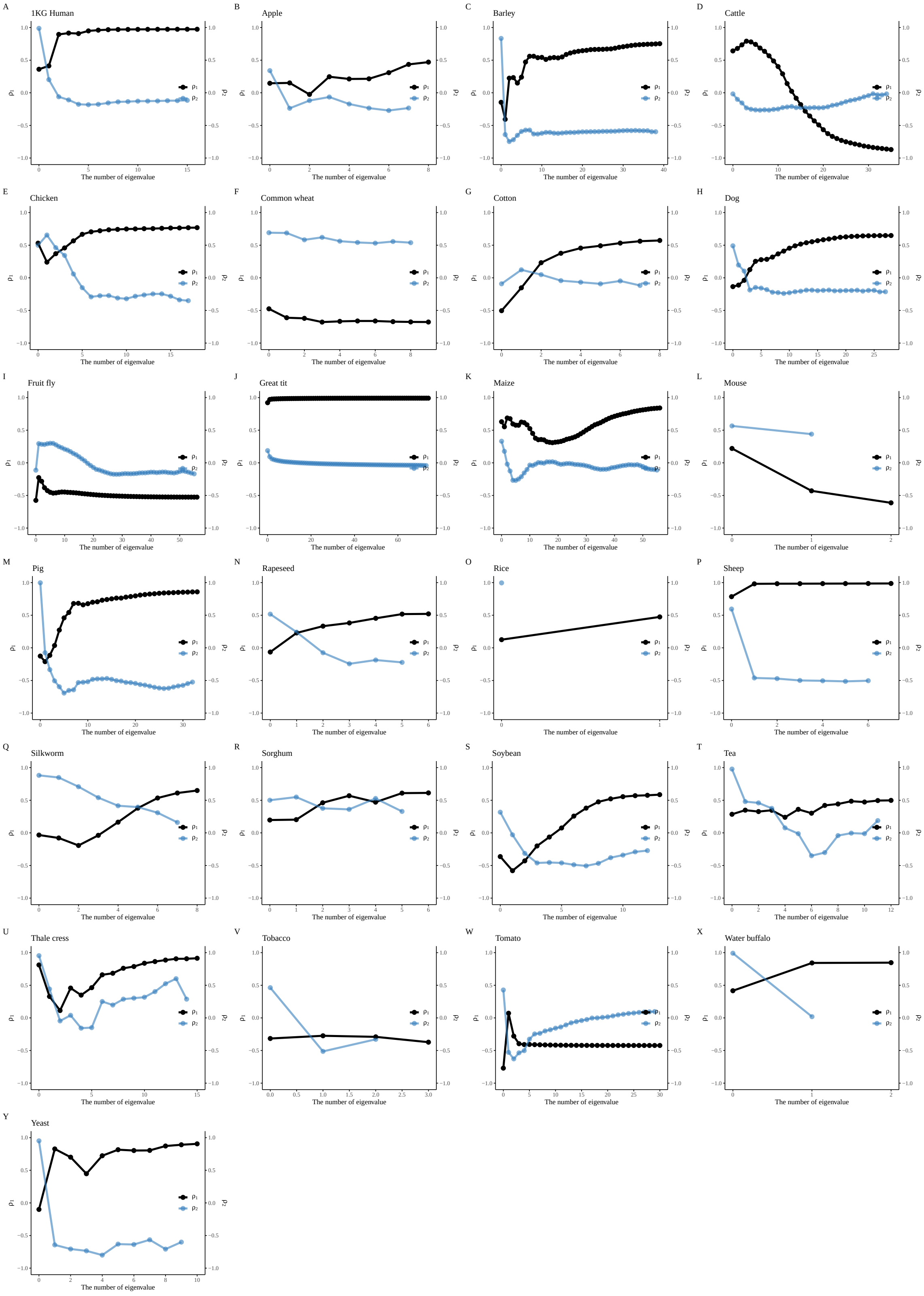


Figure S17: **The Norm I and Norm II dynamics in *RefPop*. A-Y**, the Norm I pattern and the Norm II pattern varying with the peeling of eigenvalue across **RefPop**. The black points represent the $\rho_1$ values for Norm I, while the blue points represent the $\rho_2$ values for Norm II, illustrating their reciprocal relationship.

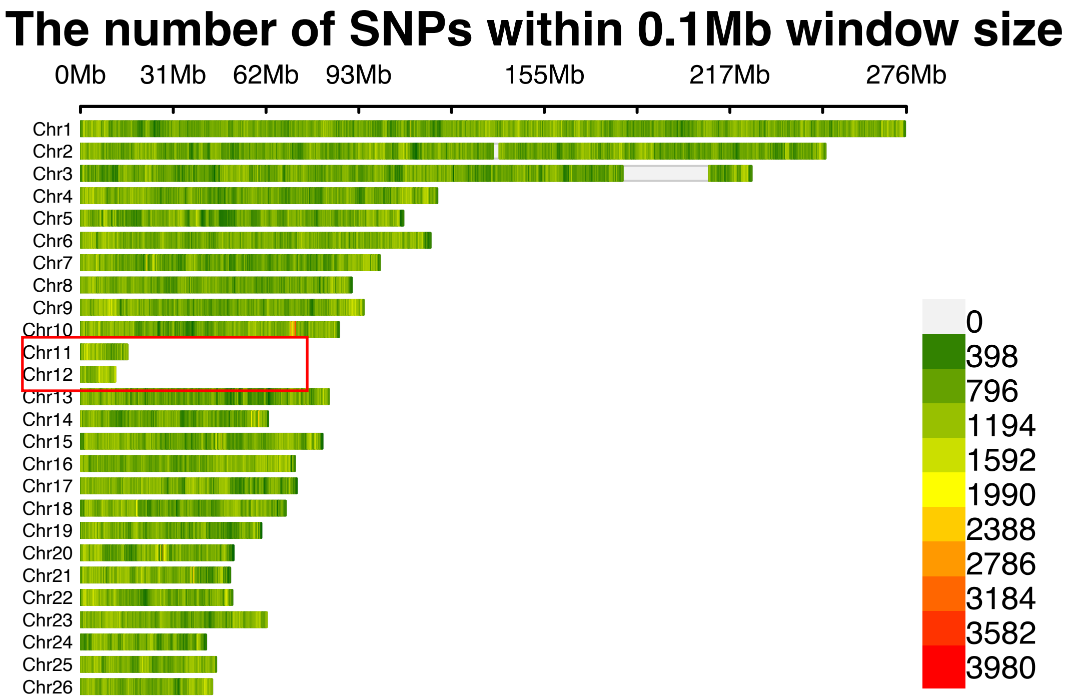


Figure S18: **SNP density plot for sheep.**

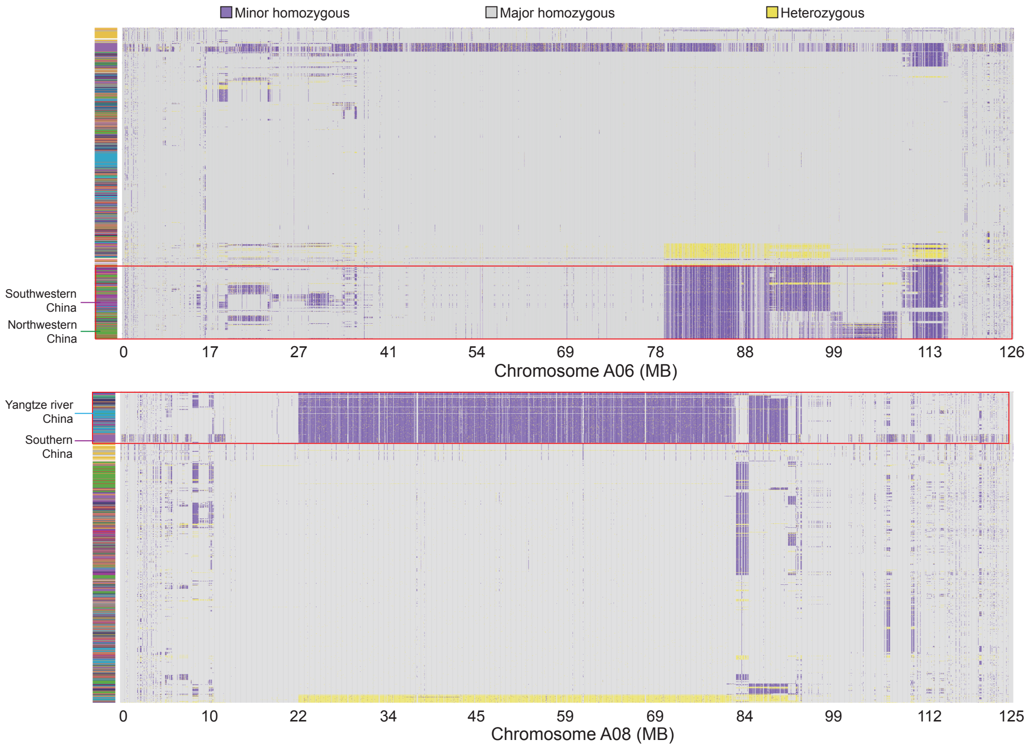


Figure S19: **Cotton haplotypes on A06 and A08.**
